# Antibiotic usage promotes the evolution of resistance against gepotidacin, a novel multi-targeting drug

**DOI:** 10.1101/495630

**Authors:** Petra Szili, Gábor Draskovits, Tamás Révész, Ferenc Bogar, Dávid Balogh, Tamás Martinek, Lejla Daruka, Réka Spohn, Bálint Márk Vásárhelyi, Márton Czikkely, Bálint Kintses, Gábor Grézal, Györgyi Ferenc, Csaba Pál, Ákos Nyerges

## Abstract

Multi-targeting antibiotics, i.e. single compounds capable to inhibit two or more bacterial targets offer a promising therapeutic strategy, but information on resistance evolution against such drugs is scarce. Gepotidacin is an antibiotic candidate that selectively inhibits both bacterial DNA gyrase and topoisomerase IV. In a susceptible organism, *Klebsiella pneumoniae*, a combination of two specific mutations in these target proteins provide an over 2000-fold increment in resistance, while individually none of these mutations affect resistance significantly. Alarmingly, gepotidacin-resistant strains are found to be as virulent as the wild-type *K. pneumoniae* strain in a murine model, and extensive cross-resistance was demonstrated between gepotidacin and ciprofloxacin, a fluoroquinolone antibiotic widely employed in clinical practice. This suggests that numerous fluoroquinolone-resistant pathogenic isolates carry mutations which would promote the evolution of clinically significant resistance against gepotidacin in the future. We conclude that prolonged antibiotic usage could select for mutations that serve as stepping-stones towards resistance against antimicrobial compounds still under development. More generally, our research indicates that even balanced multi-targeting antibiotics are prone to resistance evolution.

## Introduction

Antibiotic discovery has been driven by the need for new therapeutics that are not subject to rapid resistance development. Antibiotics selectively inhibit bacteria by targeting essential cellular processes which are absent in the human host. Most antibiotics exert their therapeutic effect by targeting a single bacterial process, and therefore they are prone to resistance development resulting from genomic mutations or the acquisition of horizontally transferred resistance genes^1^. Due to the rise of drug-resistant bacteria, the commercial success of antibiotic development is unpredictable^2,3^. For example, GlaxoSmithKline (GSK) had been focusing on the development of a novel compound (GSK2251052) that solely inhibits bacterial leucyl-tRNA synthetase. However, the clinical development of this compound was ceased, because resistance induced by mutations emerged in patients treated by GSK2251052 in its Phase 2b clinical trial^4,5^.

Antibiotic combination therapy has long been suggested as a potential strategy to delay resistance evolution^6,7^. The rationale for this theory is that drugs with different modes of action generally require different mutations in respective target genes to achieve resistance, and the simultaneous acquisition of multiple specific mutations is exceedingly rare. Although successful in many cases, antimicrobial combination therapy suffers from several limitations^1,8^. First, the efficacy of a combination therapy relies on fine-tuning the concentrations of multiple drugs *in vivo*, and this endeavor is frequently hampered by the differences in the individual drugs’ pharmacokinetic properties. Second, the prerequisite for combination therapy is the lack of mutations providing resistance to both drugs at the same time, i.e., no cross-resistance should exist between the antibiotics applied together. Third, and on a related note, the small number of available antibiotic families limits practical applications.

Multi-target antibiotic strategy is an emerging alternative that can potentially address the aforementioned challenges. There are multiple mechanisms by which antimicrobial compounds may inhibit multiple bacterial targets. In the case of hybrid drugs, two antibiotic pharmacophores with dissimilar targets are covalently linked to form one molecule. Other antibiotic compounds equipotently target two homologous, but functionally non-redundant proteins. Finally, they may target multiple, non-overlapping regions of a single bacterial protein. Currently, designing multi-target antibiotics is a major focus of the pharmaceutical industry, but until now relatively few drugs have been demonstrated to achieve a balanced inhibition of multiple microbial targets. Such exemplary antibiotics include gepotidacin^9^, NBTI 5463^10^, Trius’ C3 and C4^11^. However, due to the shortage of in-depth resistance studies, our knowledge on the tempo and mode of resistance development against multi-target antibiotics remains limited (but see^12–14^).

Gepotidacin (GSK2140944) is an exemplary candidate to study resistance evolution towards multi-targeting antibiotics (Figure 1A). Gepotidacin is a novel triazaacenaphthylene antibiotic candidate currently in phase 2b clinical trials^15,16^ and is expected to enter the clinical practice in the upcoming years^17,18^. The molecule inhibits bacterial DNA gyrase and topoisomerase IV with a novel mode of action. Using a standard frequency-of-resistance test recent studies have failed to identify resistant clones of *Neisseria gonorrhoeae* and *Escherichia coli* against this new compound^9,19^, suggesting that individual mutations cannot provide substantial resistance to gepotidacin. It is especially active against Gram-negative pathogens belonging to the *Enterobacteriaceae* family, such as *Klebsiella pneumoniae*. In fact, Carbapenem-resistant *K. pneumoniae* is resistant to nearly all commercially available antibiotics, and corresponding infections increase steadily, resulting in high rates of morbidity and mortality. According to the WHO, infections caused by multi-drug resistant *Klebsiella* strains are emerging as a top-ranked challenge in the health-care sector^20^. In this work, we demonstrate that contrary to expectations, resistance to gepotidacin evolves rapidly in *K. pneumoniae,* which has potential implications for the future clinical use of this new antibiotic candidate.

**Figure 1.**
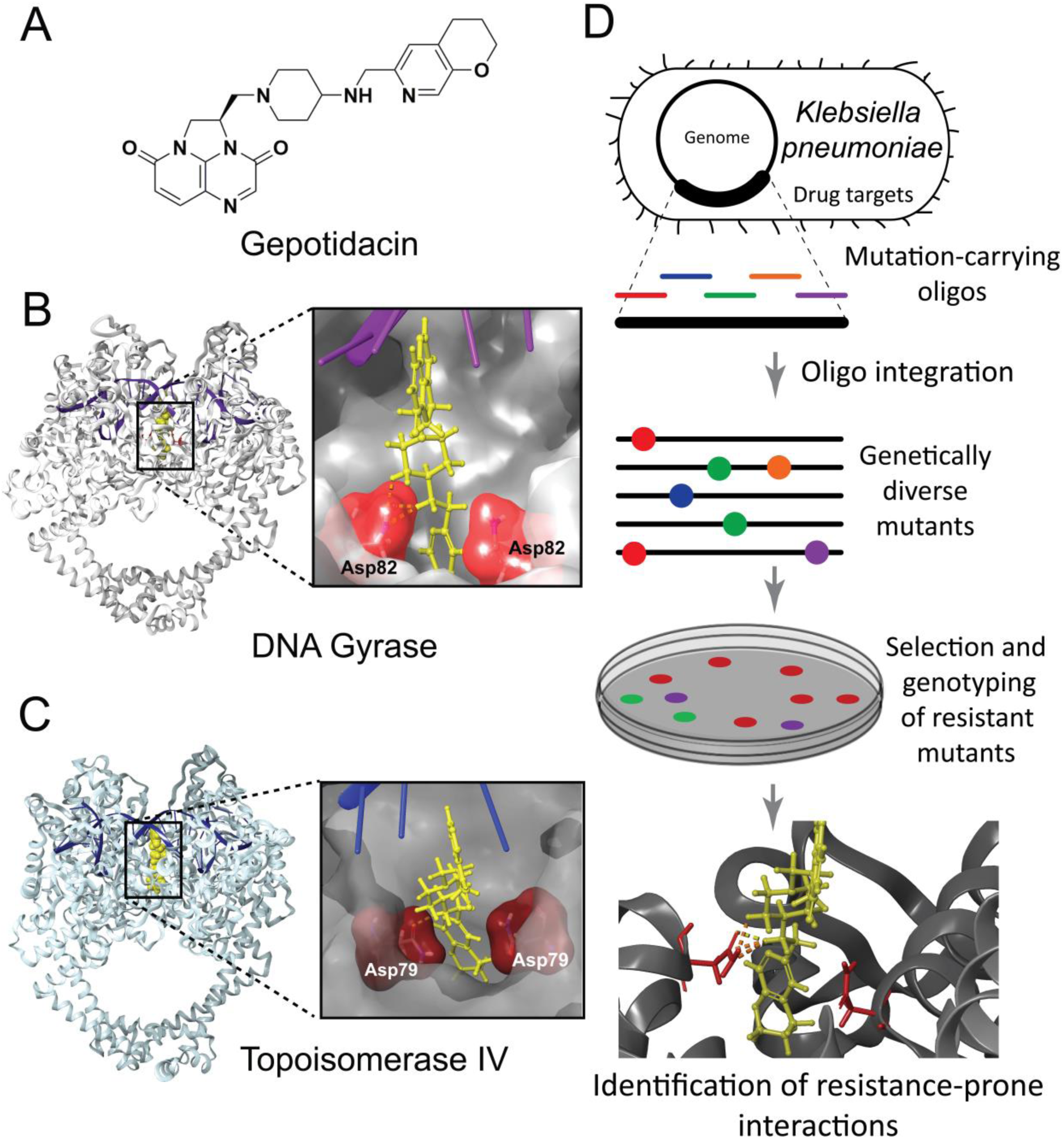
Binding site analysis of gepotidacin based on directed evolution and molecular modelling. (**A**) Gepotidacin, a novel triazaacenaphthylene antibiotic candidate inhibits DNA gyrase and topoisomerase IV in Gram-negative bacteria. (**B-C**) Ribbon representation of *E. coli*’s DNA gyrase- and topoisomerase IV in complex with gepotidacin, based on molecular dynamics simulations. Inlet shows the close-up view of gepotidacin (yellow) and its interacting residues (red) in a stick model. DNA is shown in magenta and blue. (**D**) The workflow of *in vivo* directed evolution of gepotidacin resistance in *K. pneumoniae* ATCC 10031.

## Results

### Molecular modelling with directed evolution accurately predicts the positions and impact of resistance mutations

Prior studies have demonstrated that gepotidacin selectively inhibits both bacterial DNA gyrase and topoisomerase IV by a unique mechanism^9,17^, but the exact molecular mechanisms of inhibition have not been thoroughly characterized. Therefore, we first sought to study the exact molecular interactions between gepotidacin and *Escherichia coli*’s DNA gyrase and topoisomerase IV protein complexes, using molecular modelling (see Methods). Molecular dynamics simulations capture the atomic interactions between the drug and the target protein in time. As the X-ray crystal structures of the protein targets of *E. coli* are not available, we have built homology models of DNA gyrase and topoisomerase IV based on structural templates from *Staphylococcus aureus* and *K. pneumoniae,* respectively (see Methods and Supplementary Note 1). Next, we have calculated the possible binding poses of gepotidacin using molecular docking methods. Finally, we have performed 100 ns long molecular dynamics simulations to check the stability of these binding poses, as well as to map the interaction pattern of gepotidacin and surrounding amino acid residues. The analysis has revealed that hydrophobic contacts are engulfing the pyrano-pyridine moiety of gepotidacin and the drug forms one dominant electrostatic interaction with its binding site at each target protein (Supplementary figure S1 and S2). Specifically, our model has predicted that Asp82 in the GyrA subunit of DNA gyrase and the homologous position, Asp79 of the ParC subunit of topoisomerase IV form an intermolecular salt bridge with gepotidacin (Figure 1B and C, Supplementary Note 2, and Figure S2). Therefore, mutations within this binding site are expected to provide resistance to gepotidacin. Indeed, we have provided evidence to support this hypothesis. Previously we subjected the potential target genes to a single round of DIvERGE mutagenesis in *E. coli,* and next subjected the mutant library to gepotidacin stress, followed by sequencing the resistant isolates^19^. In our current work, the analysis has been repeated in *K. pneumoniae* using nearly identical experimental settings (Figure 1D). The results of the mutagenesis assays are qualitatively the same for the two species: Asp82 within the GyrA subunit and Asp79 of the ParC subunit are found to be mutated in all the resistant clones isolated, and no further mutations have been found (Supplementary Table S3). Moreover, subsequent saturation mutagenesis of the two mutational hot-spots has recapitulated that the combination of these two specific mutations (GyrA D82N, ParC D79N) is responsible for a high-level of resistance to gepotidacin (Supplementary Table S3).

Next, to understand the molecular mechanisms behind resistance, we have computationally modelled the molecular interactions between gepotidacin and asparagine mutants at GyrA Asp82 and ParC Asp79 positions using the Molecular Mechanics/Generalized Born Surface Area (MM-GBSA)^21^ method. MM-GBSA has predicted that the observed mutations (GyrA D82N and ParC D79N) weaken the interaction between the target proteins and gepotidacin (ΔΔG_B_ = 5.44±2.92 and 4.43±1.71 kcal/mol for DNA gyrase and topoisomerase IV, respectively, based on three independent simulations). Thus, the combination of DIvERGE mutagenesis assays and molecular dynamics simulations have revealed the putative binding sites for gepotidacin in DNA gyrase and topoisomerase IV protein complexes. Furthermore, the disruption of an indispensable salt bridge between the drug and its two binding sites resulted in high-level resistance in *K. pneumoniae*.

### Exceptionally strong synergism between resistance-associated mutations

Next, we have studied the individual, as well as the combined effects of these mutations on resistance. For this purpose, the mutations were inserted individually into the genomes of *E. coli* K-12 MG1655 and *K. pneumoniae* ATCC 10031. The double-mutants were found to display an over 560-and a 2080-fold increase in gepotidacin resistance level compared to the corresponding wild-type strains of *E. coli* and *K. pneumoniae,* respectively (Figure 2, see also^22^). In contrast, the single-mutants were found to show no considerable changes in resistance level. GyrA D82N has conferred a mere 2-fold increase in resistance level in *E. coli,* while ParC D79N on its own has induced no measurable resistance. Similar results were seen for *K. pneumoniae.* These findings are consistent with prior single-step resistance selection studies that failed to recover mutants with significant resistance^9,19^.

**Figure 2.**
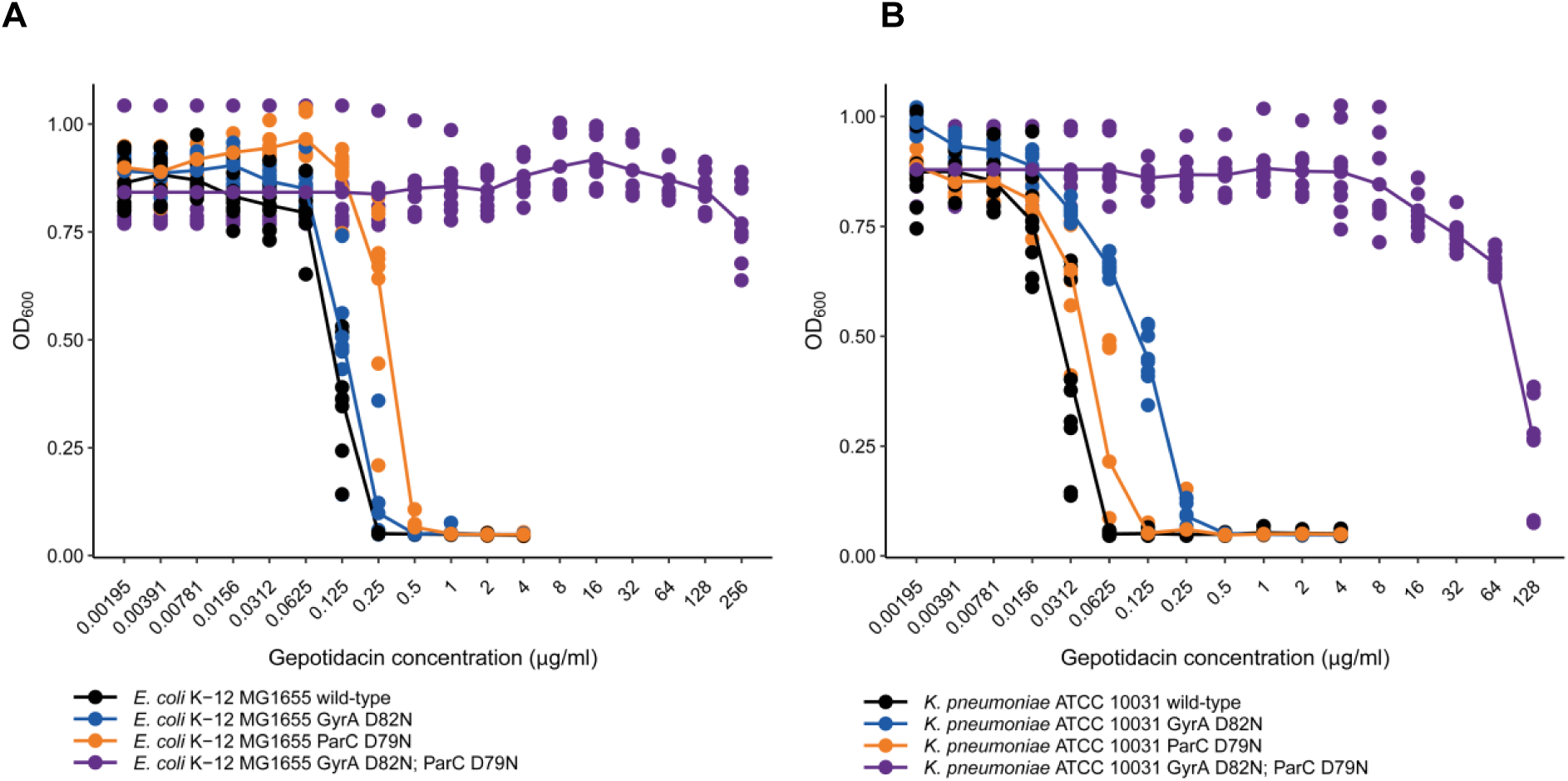
Gepotidacin dose-response curves of single-step mutations and their combination in *Escherichia coli* K-12 MG1655 (A) and *Klebsiella pneumoniae* ATCC 10031 (B). OD_600_ indicates the optical density of the bacterial culture after 18 h incubation in the presence of the corresponding drug concentration, according to the EUCAST guidelines^23^. Figure is based on 9 independent replicates; curve shows the average OD_600_ values at the corresponding drug concentration.

### Fitness cost of gepotidacin resistance is limited

The mechanisms underlying antibiotic resistance typically induce a fitness cost in the form of reduced bacterial growth rates^24^. The magnitude of fitness cost determines the long-term stability of antibiotic-resistant bacterial populations^25–27^. Thus, we have investigated the fitness effects of target-mediated resistance to gepotidacin in *K. pneumoniae*. For this aim we have studied the wild-type and mutant *K. pneumoniae* strains, the latter carrying gepotidacin or fluoroquinolone resistance-associated mutations.

The fluoroquinolone resistance-causing mutations in point are widespread in clinical isolates^28–30^ and affect the same genes (namely, *gyrA* and *parC*), therefore they provide a benchmark to estimate the fitness costs compared to the wild-type strain. Relative fitness was estimated by pairwise competition experiments between the wild-type strain and a specific mutant strain in nutrient-rich, antibiotic-free bacterial medium (MHBII) at 37°C. Using established protocols, we have found that the mutation combination of GyrA D82N and ParC D79N which confers resistance to gepotidacin significantly decreases fitness in *K. pneumoniae*. However, the fitness cost of gepotidacin resistance-associated mutations is comparable to the fitness cost of canonical, clinically relevant fluoroquinolone resistance-causing mutations. Importantly, ParC D79N mutation is found to confer no measurable fitness cost individually (Figure 3). Our findings remained the same when active human blood serum was added to the medium (see Supplementary Figure S3).

**Figure 3.**
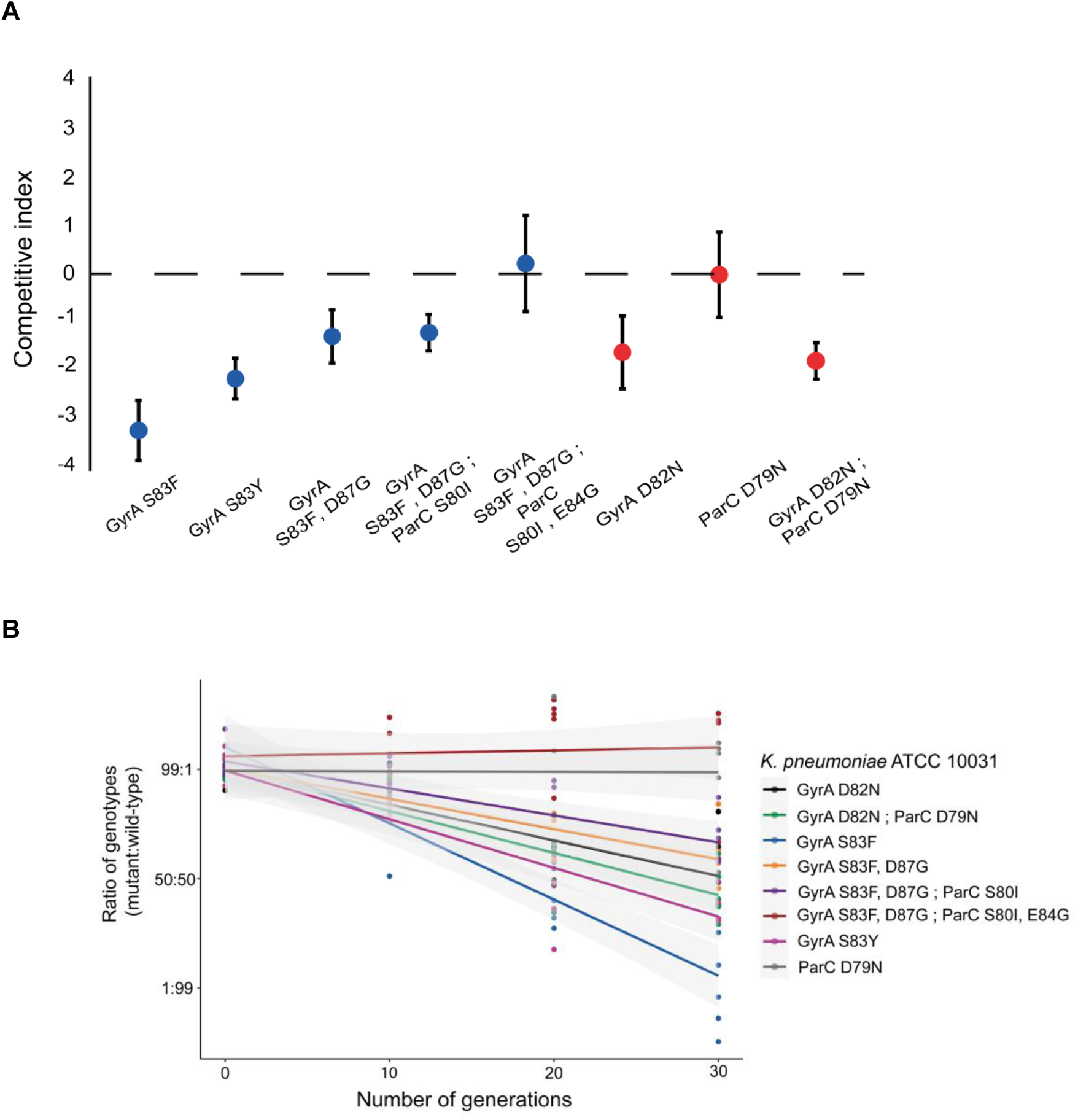
*In vitro* competition between mutant and wild-type strains of *Klebsiella pneumoniae*. (**A**) Isogenic mutants of *Klebsiella pneumoniae* ATCC 10031 carrying either a gepotidacin resistance-conferring mutation combination or its single-step constituents (red), or clinically-occurring fluoroquinolone resistance-associated mutations (blue) competed against the wild-type strain. In competition assays, a competitive index <0 indicates that the wild-type population outcompetes the mutant population, and conversely, a competition index >0 indicates that the mutant population outcompetes the wild-type population. Error bars indicate standard deviation (SD) based on five biological replicates. (**B**) Dynamics of competition between selected *K. pneumoniae* ATCC 10031 mutants and the wild-type strain as a function of bacterial generation number. The ratio of genotypes was defined as the ratio of the given mutant compared to the wild-type strain within the cell population. The gray zone indicates 95% confidence interval based on five biological replicates.

### Gepotidacin-resistant mutants display no changes in virulence in a murine infection model

Based on the above results, we next investigated whether gepotidacin-resistant mutants display reduced virulence *in vivo*. As drug-resistant *K. pneumoniae* is frequently responsible for wound and systemic infections, we have studied a murine thigh wound infection model^31,32^. Specifically, we have examined the wound colonization capacity of the wild-type strain, as well as that of representative gepotidacin- and ciprofloxacin-resistant *K. pneumoniae* mutants. For this purpose, the wild-type strain, as well as each isogenic mutant strain was inoculated into the thigh tissue of female ICR mice (*n* = 5). After 26 hours, bacterial colonization was determined by plating thigh tissue homogenates to MHBII agar plates. No significant decrease in *in vivo* virulence was observed for any of the resistant mutants compared to the wild-type strain (Figure 4).

**Figure 4.**
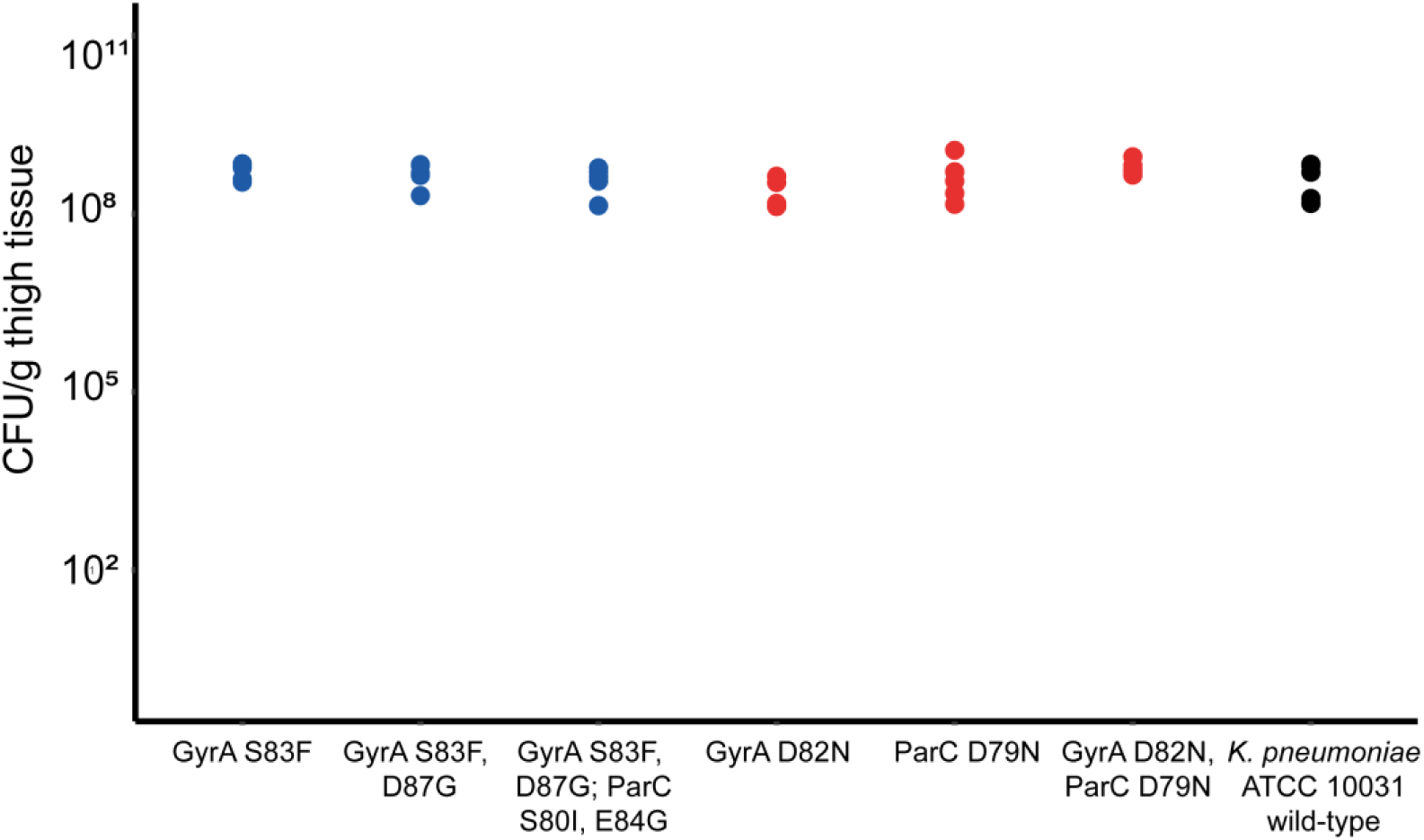
Virulence of *Klebsiella pneumoniae* mutants and the wild-type strain in a murine thigh infection model. Bacterial burden in infected thigh tissues after 26 hours of infection caused by wild-type *Klebsiella pneumoniae* ATCC 10031 (black) or isogenic mutants carrying either gepotidacin resistance-causing mutations or its single-step constituents (red), or clinically-occurring mutations associated with fluoroquinolone resistance (blue). The bacterial burden was assayed in Colony Forming Units (CFU)/g of tissue (*n* = 5 animals per data point, see Methods for details). No mutants were found to display a significant difference in virulence compared to the wild-type strain (*t*-test, P >0.05 for all pairwise comparisons).

### Cross-resistance between gepotidacin and ciprofloxacin

Taken together, the above results demonstrate that the combination of two mutations in the genes encoding gepotidactin’s target proteins confer an over 2000-fold increase in gepotidacin-resistance in *K. pneumoniae.* Alarmingly, these mutations do not seriously affect bacterial fitness and virulence *in vivo*.

Several prior works demonstrate that certain resistance mutations are present in bacterial populations long before being exposed to an antibiotic in point. Thus, we hypothesized that prolonged use of other antibiotics might select for mutations that serve as stepping-stones towards gepotidacin-resistance. The best candidate antibiotic family is fluoroquinolones, as they are widely employed in clinical practice, and similarly to gepotidacin, they target the gyrase/topoisomerase protein complexes, albeit with a notably different molecular mechanism^18,33^. Moreover, the putative binding sites of gepotidacin and fluoroquinolones on the GyrA protein are adjacent to each other, separated by a single amino acid only (see figure 1B, and^34–36^). Despite the functional similarities between these drug classes, a prior paper reported that fluoroquinolone-resistant clinical isolates displayed no cross-resistance to gepotidacin^18^. To re-investigate this issue, we have focused on ciprofloxacin, a widely employed and well-characterized fluoroquinolone drug.

We have found that the GyrA D82N single-mutant strain displays an over 16-fold increase in ciprofloxacin-resistance compared to the wild-type *K. pneumoniae* ATCC 10031 strain. This finding is even more remarkable if we consider that the same mutation confers only a 2-fold increase in the level of resistance to gepotidacin. As it might be expected, the GyrA D82N and ParC D79N double-mutant also displays significant resistance to ciprofloxacin (Figure 5).

**Figure 5.**
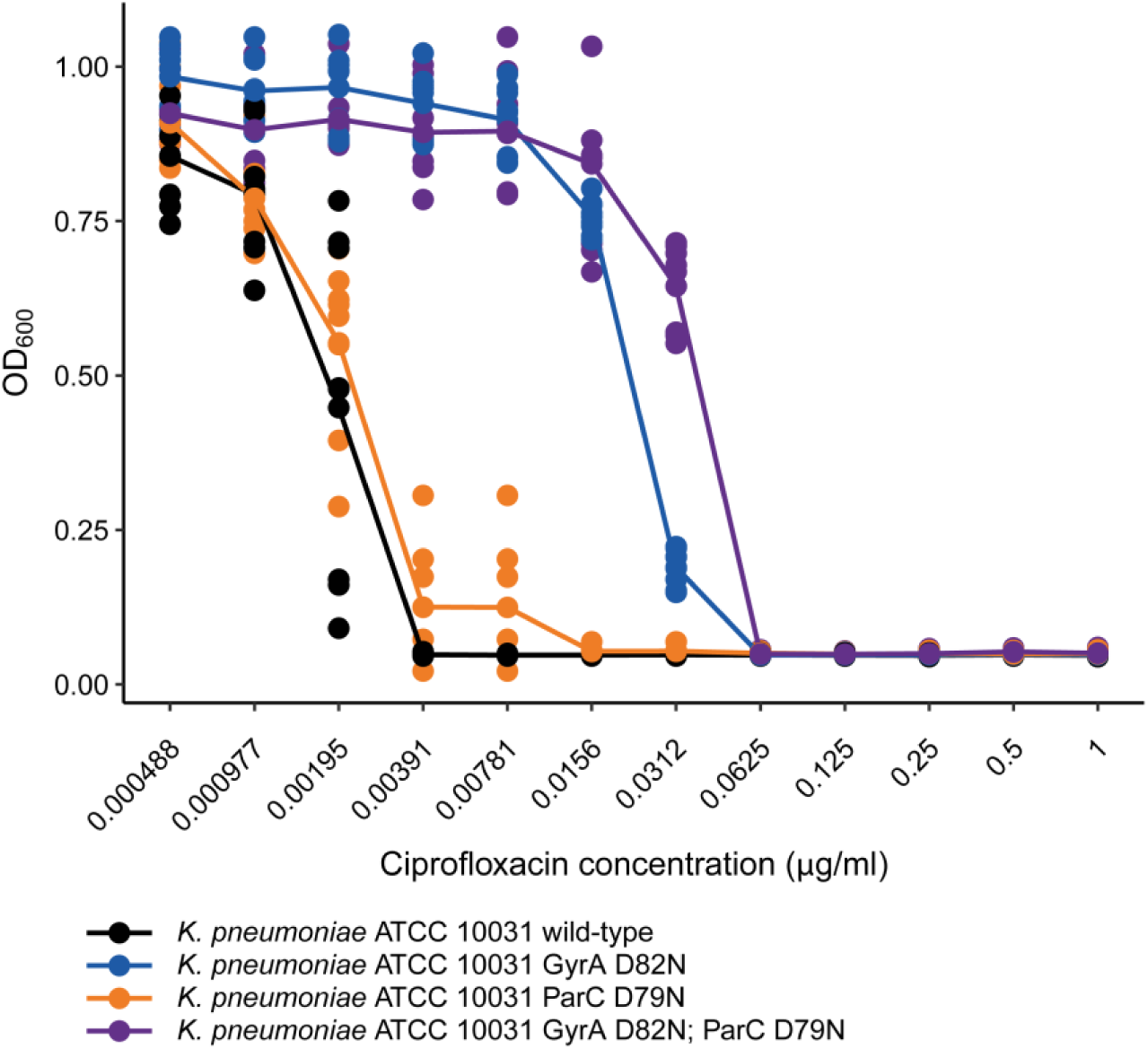
Ciprofloxacin dose-response curves of gepotidacin resistance-associated single-step mutations and their combination in *Klebsiella pneumoniae*. OD_600_ indicates the optical density of the bacterial culture after 18 h incubation in the presence of the corresponding drug concentration, according to the EUCAST guidelines^23^. Figure is based on 9 independent biological replicates. The curve shows the average OD_600_ values at the corresponding drug concentration.

Our findings raise the possibility that the D82N mutation of GyrA might be present in fluoroquinolone-resistant clinical isolates, rendering the subsequent emergence of gepotidacin-resistant double-mutants feasible. Indeed, GyrA D82N or ParC D79N mutants have been detected in a wide range of fluoroquinolone-resistant clinical isolates of *E. coli*^37^, *Salmonella sp.*^38–41^, *Neisseria gonorrhoeae*^42,43^, *Mycoplasma genitalium*^44^ (Table 1) and also in laboratory experiments^45–48^. In line with this hypothesis, we have demonstrated that the mutation combination of GyrA D82N, existing in clinical isolates, and an additional D79N mutation at ParC causes a more than 128-fold increase in gepotidacin-resistance in *S. enterica* (Figure 6). These results indicate that several clinically occurring human pathogens require only a single extra mutation to evolve high-level resistance to gepotidacin.

**Table 1.**
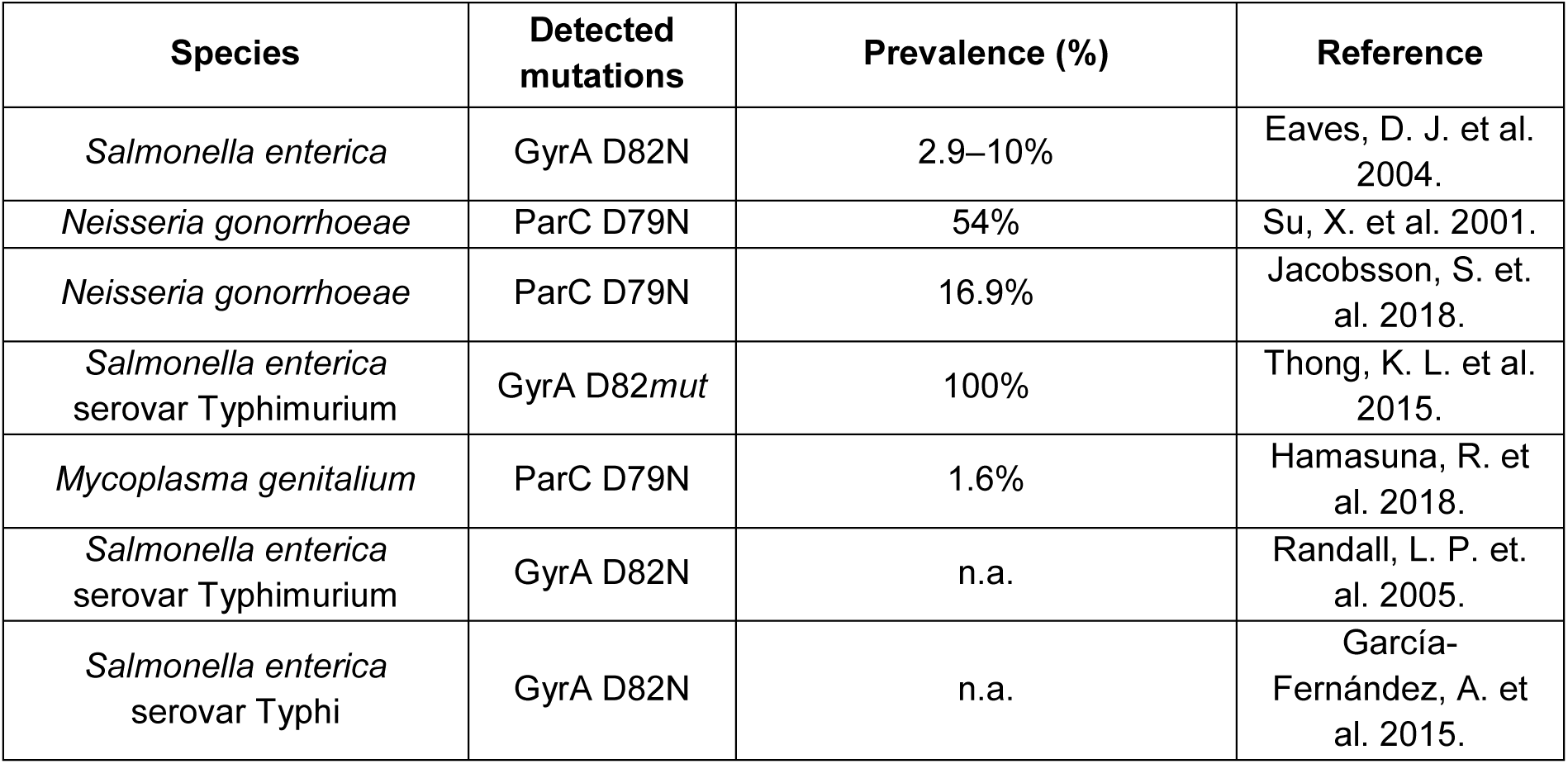
Fluoroquinolone resistance-associated mutations and their prevalence in human clinical pathogens. Amino acid positions are marked according to their corresponding residue in *Escherichia coli* K-12 MG1655. N.a denotes observed mutation (prevalence = single case), *mut* denotes mutation at a given position.

**Figure 6.**
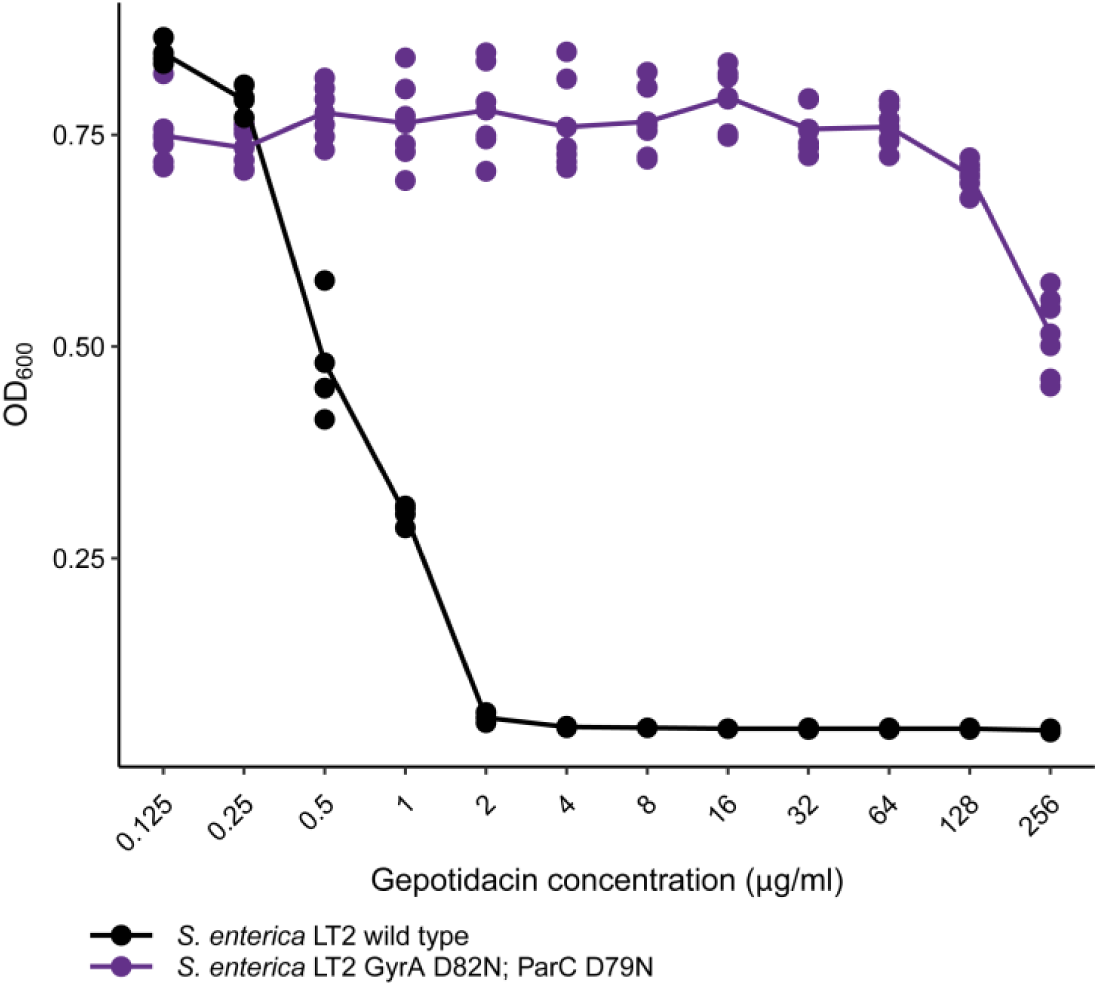
Gepotidacin dose-response curve of the gepotidacin resistance-associated mutation combination in *Salmonella enterica*. OD_600_ indicates the optical density of the bacterial culture after 18 h incubation in the presence of the corresponding drug concentration, according to the EUCAST guidelines^23^. Figure is based on 6 independent biological replicates. The curve shows the average OD_600_ values at the corresponding drug concentration.

Given the high frequency of mutations associated with potential gepotidacin resistance in clinical isolates, next we have systematically investigated the prevalence of the GyrA D82N and ParC D79N mutations in currently available sequence databases (as of 9 October 2018). Systematic sequence search has revealed that both mutations occur in a wide-range of Gram-negative and Gram-positive bacteria (Supplementary Figure S4 and Supplementary File 1), including species belonging to the *Mycoplasma, Clostridium, Citrobacter, Streptococcus,* and *Neisseria* genera. *Neisseria* and *Streptococcus* are especially noteworthy, as infections caused by these genera are reported to be the targets of gepotidacin in recent clinical trials^15,16,49^.

### Ciprofloxacin stress selects for gepotidacin resistance in the laboratory

To establish that ciprofloxacin promotes gepotidacin resistance, we have initiated laboratory evolution under ciprofloxacin stress. We followed established protocols with minor modifications to evolve bacterial populations under controlled laboratory conditions^50–52^. The protocol aims to maximize the level of drug resistance in the evolving populations that develop in a fixed time period, being approximately 116 generations in our case. For our laboratory evolutionary experiments, we have chosen the wild-type *K. pneumoniae* ATCC 10031, as well as the wild-type and Δ*mutS* variants of *E. coli* K-12 MG1655. Six parallel evolving populations of every strain were exposed to gradually increasing concentrations of ciprofloxacin. Δ*mutS* displays around 100-fold increase in mutation rate, and as being a hypermutator, it adapts especially rapidly to various antibiotic stresses. In line with previous clinical and laboratory studies^33,53^, ciprofloxacin resistance was seen to emerge quickly in our experiments (Figure 7). Despite the short time-frame, all *E. coli* and *K. pneumoniae* populations reached ciprofloxacin-resistance levels equal to or above the EUCAST clinical breakpoint^54^ (Figure 7). As expected, the evolution of clinically significant resistance was especially rapid in mutator populations.

**Figure 7.**
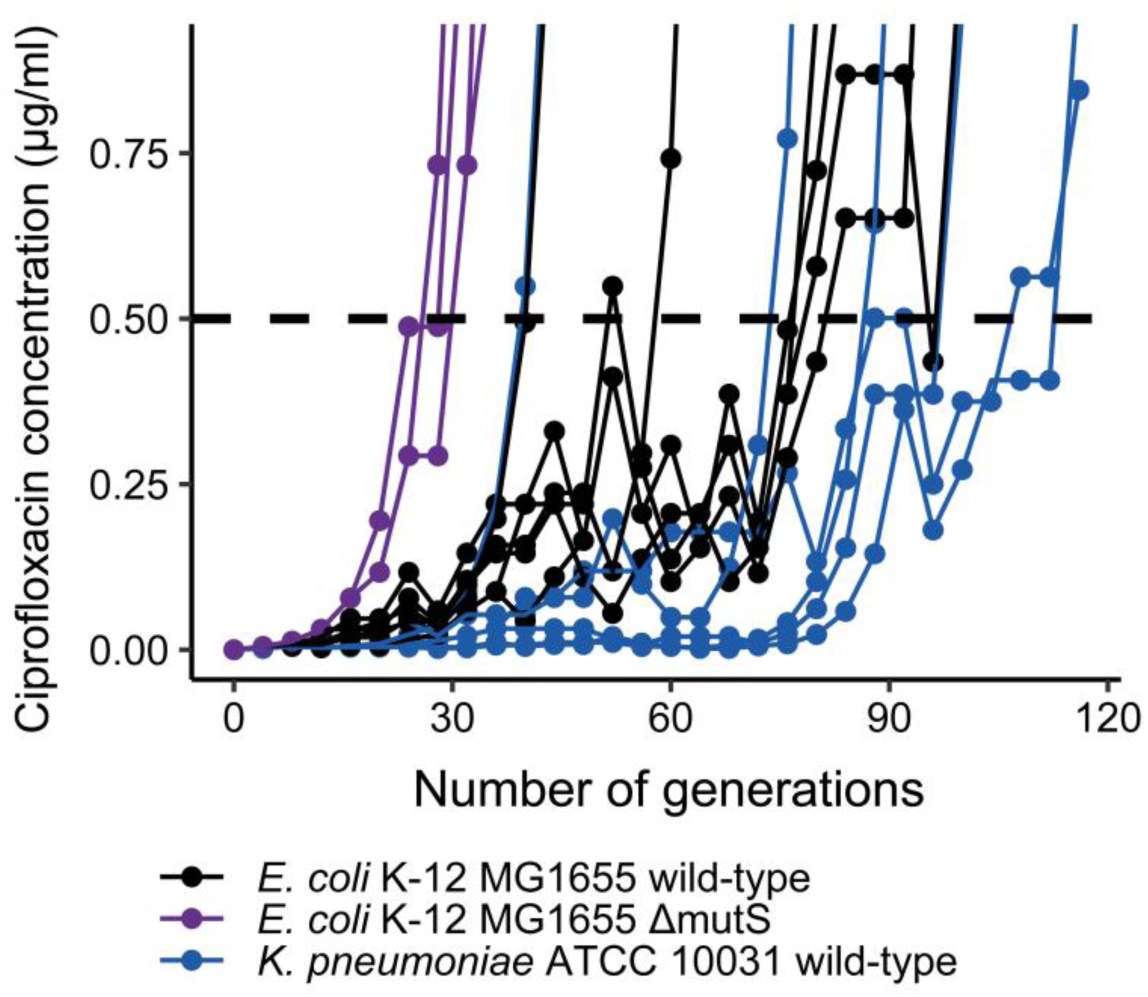
Adaptive laboratory evolution of *Klebsiella pneumoniae* and *Escherichia coli* under ciprofloxacin stress. Figure shows the antibiotic concentrations at which *K. pneumoniae* ATCC 10031, *as well as E. coli* K-12 MG1655 wild-type and Δ*mutS* hypermutator strains were able to grow under increasing ciprofloxacin stress as a function of time (number of cell generations). Dashed line represents the clinical breakpoint of ciprofloxacin resistance according to EUCAST^54^.

Strikingly, ciprofloxacin-resistant lineages of both species have been found to display an over 64 to 512-fold increment in gepotidacin-resistance level compared to the corresponding wild-type strains (Figure 8). Some, but not all of these gepotidacin-resistant lineages carried mutations at GyrA D82 and ParC D79 (Supplementary Table S2), suggesting that multiple other mutations may also select for gepotidacin resistance. Overall, these results strongly suggest that long-term exposure to ciprofloxacin stress promotes resistance to gepotidacin.

**Figure 8.**
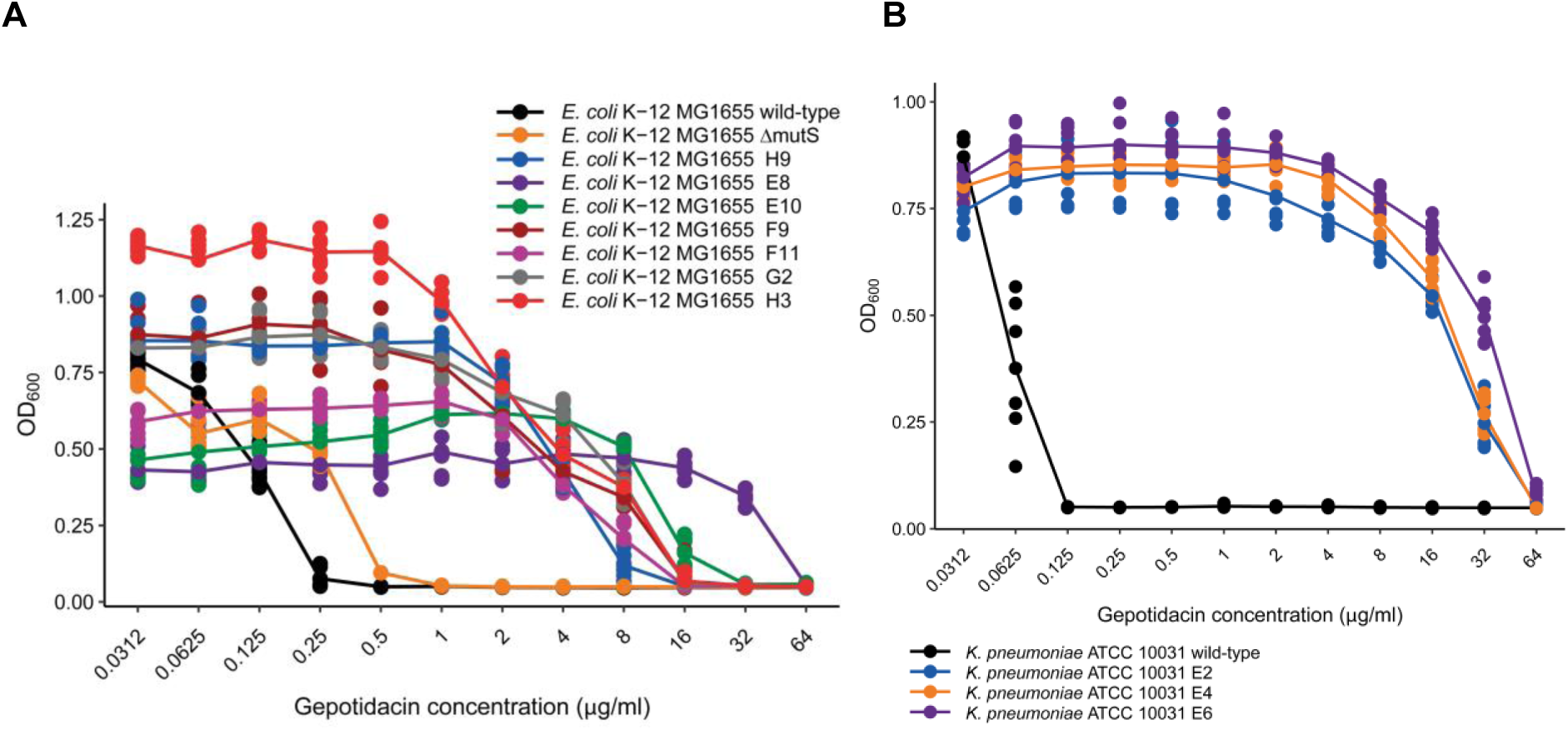
Gepotidacin dose-response curves of ciprofloxacin-adapted *E. coli* (A) and *K. pneumoniae* (B) isolates. OD_600_ indicates the optical density of the bacterial culture after 18 h incubation in the presence of the corresponding drug concentration, according to the EUCAST guidelines. Figure shows gepotidacin dose-response of independently evolved, ciprofloxacin-resistant *E. coli* K-12 MG1655 and *K. pneumoniae* ATCC 10031 isolates and their corresponding parental strains. Figure is based on 6 independent biological replicates. The curve shows the average OD_600_ values at the corresponding drug concentration. For the genotypic analysis of each isolate, see Supplementary Table S2.

## Discussion

Gepotidacin is a novel bacterial topoisomerase inhibitor (NBTI) antibiotic candidate that simultaneously inhibits both bacterial gyrase and topoisomerase IV with a novel mechanism of action. Therefore, it is an excellent candidate to study the evolution of resistance against dual-targeting antimicrobials. As it is still under clinical development, it is highly relevant to predict the tempo and mechanisms of resistance development against this new drug candidate.

As a good benchmark for mono-targeting drugs, both mutation rate and population size should be high enough to test at least three mutations across 99.9% of nucleotide sites in the genome, requiring that 2×10^11^ wild-type cells are assayed is in a frequency-of-resistance assay^50^. For multi-targeting drugs, this mutational space scales exponentially with the number of targets. Thus, for a drug with two bacterial targets, screening over 10^22^ wild-type *E. coli* cells would be required to identify mutation combinations that may confer resistance. Strains with an elevated genomic mutation rate (hypermutators) could potentially reduce this astronomical number, but testing over 10^18^ mutator bacteria still remains infeasible^55,56^. Recent genome-engineering technologies, however, offer promising alternatives to explore resistance-conferring mutations in a more systematic manner^19,57–60^. One of these methods termed directed evolution with random genomic mutations (DIvERGE)^19^ allows an up-to a million-fold increase in mutation rate along the full length of bacterial genes encoding drug targets. Thus, it is capable to rapidly explore specific mutational combinations that confer resistance to antibiotics.

Using DIvERGE, we have primarily focused on studying the evolution of resistance to gepotidacin in *Klebsiella pneumoniae* (Figure 1D). Currently, the rise of drug-resistant *Klebsiella* infections is especially alarming, as this pathogen is frequently responsible for life-threatening diseases, including pneumonia, sepsis and wound infections. Isolates that are resistant to all marketed antibiotics have already been detected^28^. As gepotidacin shows excellent potency against Gram-negative bacteria^17,18,61^, it could offer a last resort drug to treat multidrug-resistant *Klebsiella* infections.

However, we have demonstrated that two specific mutations in the genes encoding gepotidacin’s targets can provide a very high level of resistance in multiple enterobacterial species^22^ (see also Figure 2 and 6). These two mutations (GyrA D82N and ParC D79N) overlap with the predicted binding sites of the drug and show extreme synergism. By abolishing indispensable drug-target interactions, they confer an over 2000-fold increment in resistance level, but individually they have only limited effects on gepotidacin-resistance. Alarmingly, the double-mutant *K. pneumoniae* strain is as virulent as the wild-type in a mouse infection model, suggesting that these mutations might have clinical significance. Indeed, gepotidacin-resistant *Neisseria gonorrhoeae* strains isolated in a human clinical phase 2b study were shown to carry a ParC D86N mutation^49^ that corresponds to the homologous ParC D79N mutation in *E. coli*.

According to what we term the “stepping-stone” hypothesis, prolonged clinical deployment of certain antibiotics may select for variants with an elevated potential to evolve resistance to new antimicrobial agents. For example, in *Staphylococcus aureus* the genetic alteration responsible for methicillin-resistance had emerged due to the selective pressure of first-generation beta-lactams, such as penicillin, years before methicillin was first applied in clinical practice^62^. Furthermore, the byproducts of drug degradation were also shown to promote resistance development against clinically applied antibacterials^63^.

To investigate the feasibility of the “stepping-stone” hypothesis, here we have focused on ciprofloxacin, a widely employed and well-characterized fluoroquinolone antibiotic^64^. Similarly to gepotidacin, mutations in GyrA and ParC contribute to resistance to ciprofloxacin. Strikingly, gepotidacin resistance-linked mutations (i.e., GyrA D82N, ParC D79N) increase resistance to ciprofloxacin and have been detected in clinical isolates of ciprofloxacin-resistant bacteria, including strains of *N. gonorrhoeae, Streptococcus sp.,* and *S. enterica*. Therefore, these isolates are expected to require only one extra mutational step to develop a high level of gepotidacin resistance. In line with this hypothesis, gepotidacin failed to be effective in patients carrying a ParC D86N mutant strain of *N. gonorrhoeae* in a Phase 2b clinical trial (n.b. ParC D86N is homologous to ParC D79N in *E. coli*).

The “stepping-stone” hypothesis may have general implications. For example, zoliflodacin (ETX0914), a novel bacterial topoisomerase inhibitor has just completed a human phase 2 clinical trial^65^. It shows promising activity against multidrug-resistant infections, including *N. gonorrhoeae*^66^. However, in this species mutations in GyrB have been reported to confer zoliflodacin-resistance^67^, and one of these zoliflodacin-resistant mutants (D429N) has already been detected in clinical populations^68^. As the homologous GyrB mutation (D426N) in *E. coli* confers resistance to fluoroquinolones^22^, fluoroquinolone agents might have incidentally selected for reduced susceptibility to zoliflodacin as well. Consistent with this hypothesis, naturally occurring zoliflodacin-resistant variants were also found to be highly resistant to fluoroquinolones^68^.

In summary, our work demonstrates that despite a balanced *in vivo* targeting of multiple proteins, high-level resistance can rapidly emerge to antibiotics when the drug molecule’s inhibitory effect depends merely on interactions with a few, indispensable amino acids. Moreover, based on adaptive laboratory evolution and clinical data, we propose that target gene mutations conferring resistance to fluoroquinolones can facilitate resistance evolution to novel topoisomerase-targeting antimicrobials, including gepotidacin. Therefore, existing resistant bacteria could serve as an immediate source of novel resistance mechanisms.

As a more general concept, our work indicates that even drugs executing a balanced multi-targeting of bacterial proteins are prone to resistance, and multi-targeting itself does not preclude the appearance of high-level resistance to antimicrobials. Regarding that a quarter of the antibiotics currently under clinical development target bacterial topoisomerases^69^, further research is warranted to test this scenario.

## Acknowledgments

We thank Donald L. Court (National Cancer Institute, USA) for providing *Salmonella enterica* LT2. We also wish to thank Lynn Miesel, Andrea Tóth, Lucy Chia, and Tamás Kukli for their support. This work was supported by grants from the European Research Council H2020-ERC-2014-CoG 648364 ‘Resistance Evolution’ (to C.P.), the Wellcome Trust (to C.P.), and GINOP (MolMedEx TUMORDNS) GINOP-2.3.2-15-2016-00020, GINOP (EVOMER) GINOP-2.3.2-15-2016-00014 (to C.P.), EFOP 3.6.3-VEKOP-16-2017-00009 (to P. Sz. and T.R.) and UNKP-18-3 New National Excellence Program of the Ministry of Human Capacities (to P. Sz.); the ‘Lendület’ Program of the Hungarian Academy of Sciences (to C.P.), an NKFIH grant K120220 (to B.K.), and a PhD fellowship from the Boehringer Ingelheim Fonds (to Á.N.). B.K. was supported by the UNKP-18-4 New National Excellence Program of the Ministry of Human Capacities, the János Bolyai Research Scholarship of the Hungarian Academy of Sciences, and M.C. was supported by the Szeged Scientists Academy under the sponsorship of the Hungarian Ministry of Human Capacities (EMMI: 13725-2/2018/INTFIN). The authors thank Dora Bokor PharmD for proofreading the manuscript and acknowledge KIFÜ for awarding us access to resource based in Hungary at Debrecen.

## Authors’ contributions

The research was conceived and supervised by Á.N., and C.P.; T.R., Á.N., G.D., P.S., F.B., and C.P. designed the experiments and interpreted experimental data. G.D., P.S., Á.N., G.F., M.S., T.R., D.B., L.D., R.S., T.M., M. C., G.G., B.K., V.B.M. performed the experiments or calculations.

## Competing financial interests

The authors declare competing financial interest. Á.N., B.K., and C.P. have filed a patent application (PCT/EP2017/082574) related to DIvERGE.

## Supplementary Information

**Supplementary Note 1. Homology model construction for *E. coli* DNA gyrase and topoisomerase IV.** To construct homology models of *E. coli* DNA gyrase and topoisomerase IV, homologous bacterial protein sequences were downloaded from UniProt (https://www.uniprot.org/) (see also Methods). Sequence alignments were carried out using the Schrödinger Program Suite^70^ with BLOSUM62 substitution matrix, with -10.0 as gap opening and -1.0 as gap extension penalty. **Supplementary Table S1** lists sequence identities and homologies of target-template sequence pairs used in our homology model construction.

**Supplementary Table S1.**
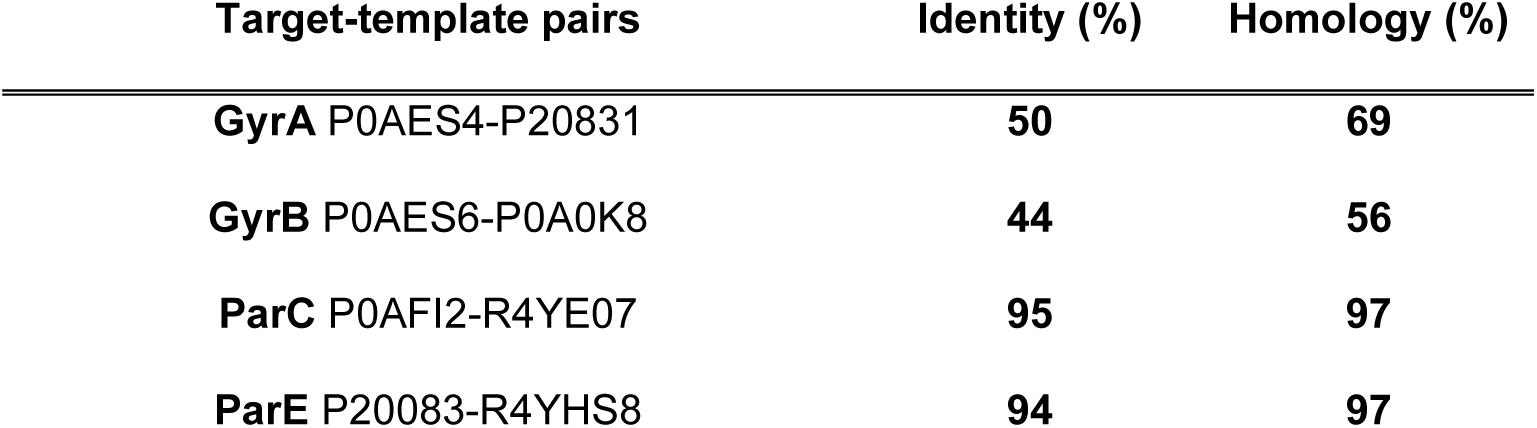
Sequence identity and homology of target sequences GyrA(P0AES4), GyrB(P0AES6) chains of DNA gyrase and ParC(P0AFI2), ParE(P20083) chains of topoisomerase IV of *E. coli* K-12, compared to their template structures from *Staphylococcus aureus* (GyrA(P20831), GyrB(P0A0K8)) and *K. pneumoniae* (ParC(R4YE07), ParE(R4YHS8)), respectively.

| Target-template pairs | Identity (%) | Homology (%) |
| --- | --- | --- |
| **GyrA** P0AES4-P20831 | **50** | **69** |
| **GyrB** P0AES6-P0A0K8 | **44** | **56** |
| **ParC** P0AFI2-R4YE07 | **95** | **97** |
| **ParE** P20083-R4YHS8 | **94** | **97** |

**Supplementary Note 2. Two alternative binding poses of gepotidacin at *E. coli* DNA gyrase and topoisomerase IV.** The binding sites for gepotidacin at both DNA gyrase and topoisomerase IV are located at the interface of the GyrA and ParC subunits, respectively (Figure 1). These two subunits together form the mainly hydrophobic binding pocket of gepotidacin. Based on our model, only a single strong, non-hydrophobic interaction appears between gepotidacin and the target proteins, namely a salt-bridge formed by gepotidacin with Asp82 in the GyrA subunit of DNA gyrase and Asp79 in the ParC subunit of topoisomerase IV. However, due to the symmetry of this binding cavity at both subunits (GyrA and ParC), the reconstructed binding modes contained two alternative binding poses (see Supplementary Figure S1). Both of these binding poses show an approximately 2-fold rotational symmetry in their orientation, and this way the triazaacenaphthylene ring intercalates DNA in a 180 degree-rotated orientation compared to each other. The protonated secondary nitrogen of gepotidacin also changes its dominant interaction with D82 of GyrA and D79 of ParC from one chain to the other, respectively, on both targets. The pyranopiridine ring is also turned around, but it occupies the same hydrophobic pocket. Because of the symmetrical arrangement of the two binding modes, only one of them was used in our further analyses. As a confirmation for these binding modes, the molecular dynamics simulations also proved that the initial binding positions of gepotidacin in wild-type proteins are stable: the average root mean square deviations (RMSD) of heavy atom coordinates of gepotidacin during the second half of the simulation from the initial pose were 1.24±0.27Å for the wild-type DNA gyrase, and 1.10±0.14Å for the wild-type topoisomerase IV. This means that the initial positions were somewhat changed during the simulation, but the systems could reach equilibrium positions close to the initial one with small average fluctuations.

**Supplementary Figure S1.**
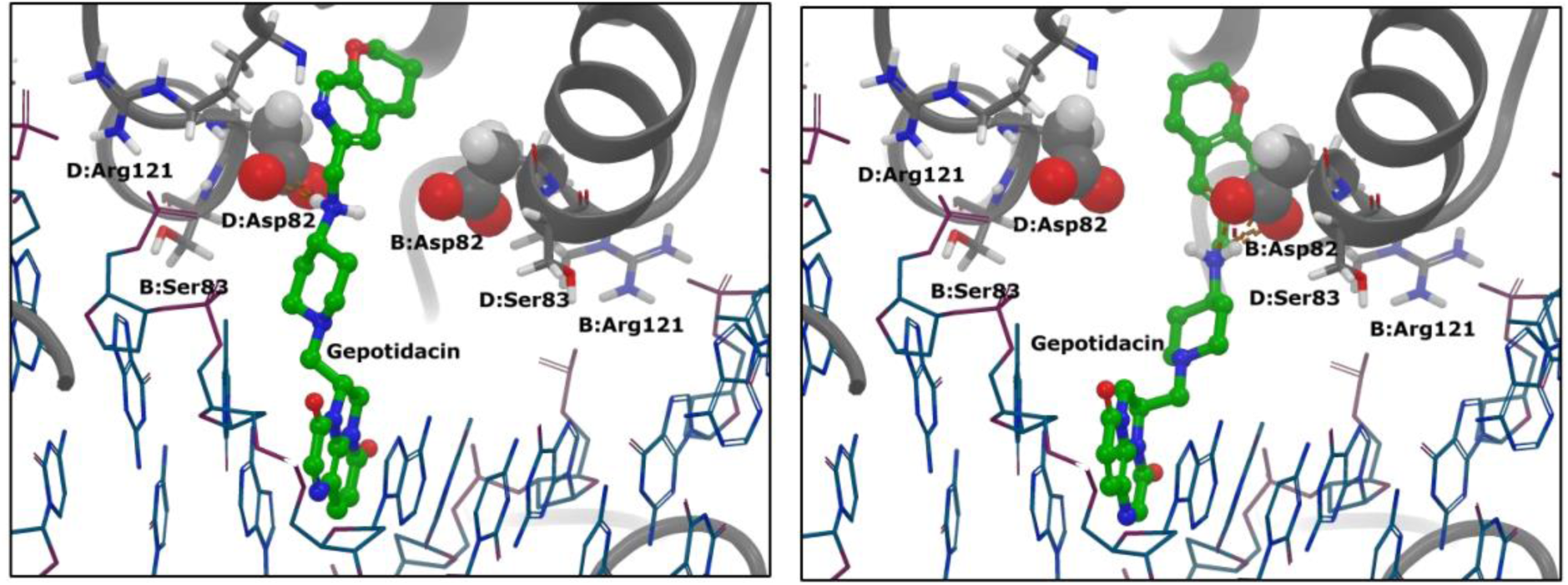
Two alternative binding poses of gepotidacin in the homology model of *E. coli* K-12 DNA gyrase. Gepotidacin appears in ball-and-stick representation, the DNA chain is shown as thin sticks, the two GyrA chains (B and D) are in ribbon representation, and Asp82 residues of GyrA are represented as balls, while Ser83 and Arg119 appear as sticks.

**Supplementary Figure S2.**
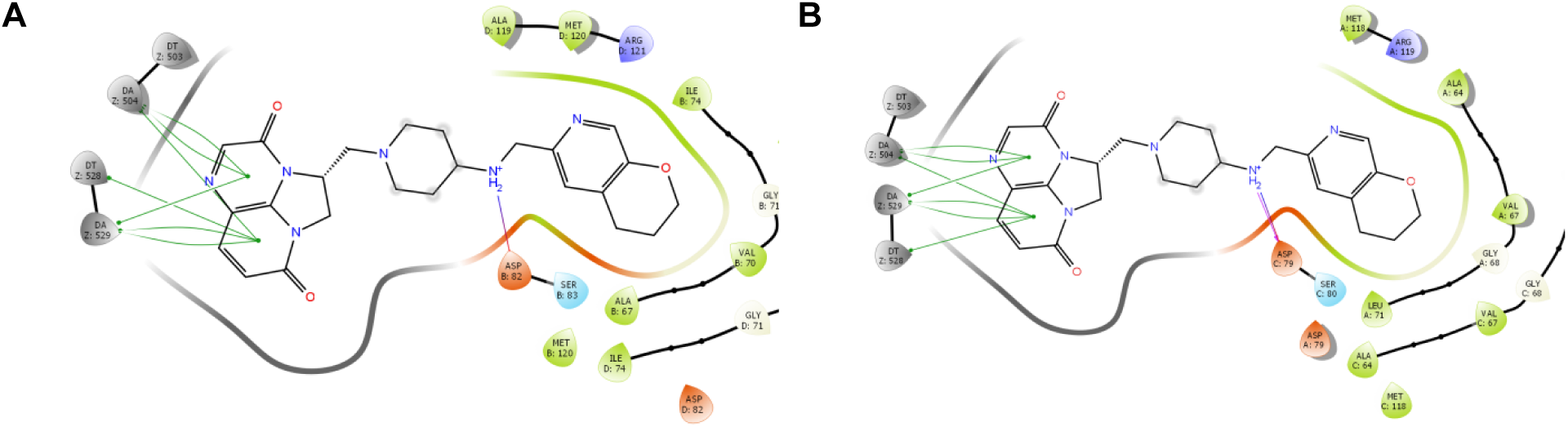
Intermolecular interactions between gepotidacin and its binding-site forming amino acids at DNA gyrase (A) and topoisomerase IV (B). Green lines indicate drug-target interactions with DNA nucleobases; purple line indicates salt bridge between gepotidacin and GyrA Asp82 (**A**), and ParC Asp79 residues (**B**) respectively. Results are based on 100 nanosecond long molecular dynamics simulations of gepotidacin-bound *E. coli* DNA gyrase and topoisomerase IV complexes. Figure displays binding-site forming amino acids and DNA nucleobases (in grey) that are closer than 4Å to gepotidacin at DNA gyrase and topoisomerase IV, respectively.

**Supplementary Figure S3.**
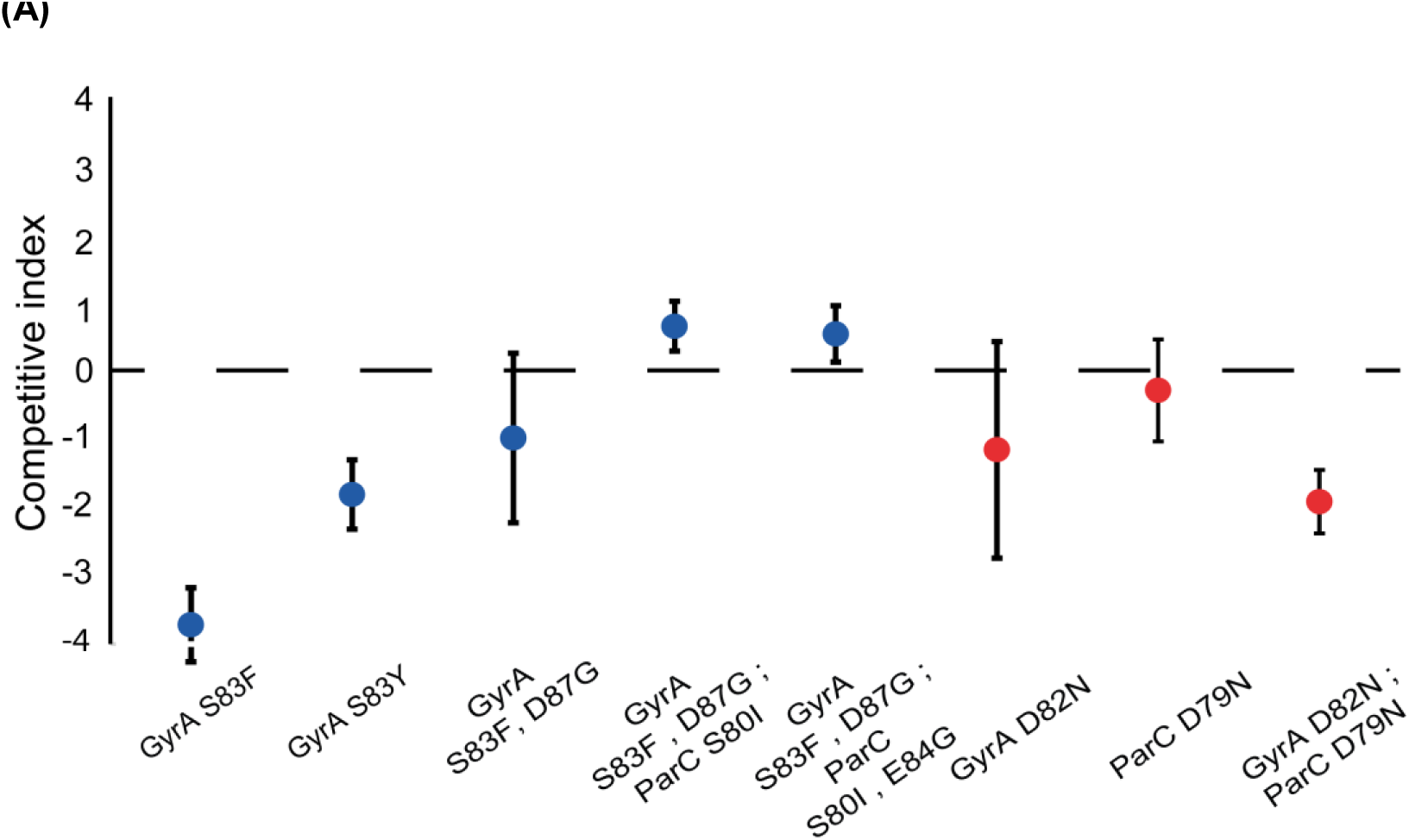
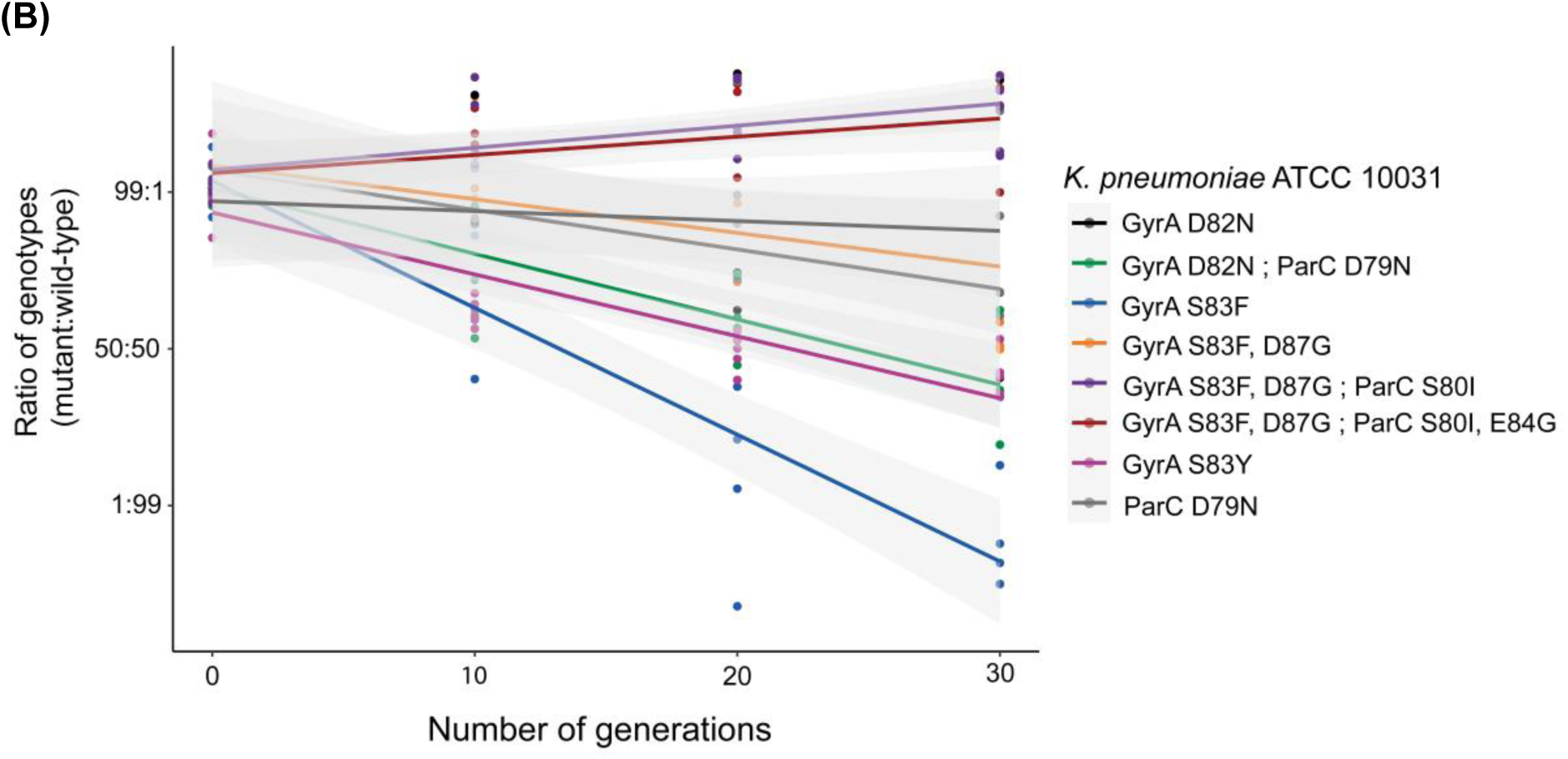
Competition assay in the presence of active human blood serum. **(A)** Applying a sub-inhibitory concentration of active human serum (i.e., 2 volume/volume%), isogenic mutants of *Klebsiella pneumoniae* ATCC 10031 carrying either the gepotidacin resistance-conferring mutation combination, its single-step constituents (red), or clinically occurring, fluoroquinolone resistance-linked mutations (blue) were competed against the wild-type strain. Importantly, all strains were equally susceptible to human blood serum (MIC = 6.7 volume/volume%). A competitive index <0 indicates that the wild-type population outcompetes the mutant population, and conversely, a competition index >0 represents that the wild-type population is outcompeted by the mutant. Error bars indicate standard deviation (SD) based on five replicate measurements. **(B)** Dynamics of competition between selected *K. pneumoniae* ATCC 10031 genotypes and the wild-type bacteria in the function of bacterial cell-generation number, in the presence of active human blood serum. The ratio of genotype is defined as the ratio of the given mutant compared to the wild-type. Gray zone marks 95% confidence interval based on five replicates.

**Supplementary Table S2.** **Mutations at GyrA and ParC in the ciprofloxacin-adapted lines after laboratory evolution experiments.** Wt represents wild-type strains and Δ*mutS* represents the deletion of *mutS*, yielding a hypermutator phenotype. Generation number represents the cell-generation number at which the given isolate was obtained from the adaptive laboratory experiment.

| Organism | Clone ID | Antibiotic selection | Generation number at isolation | Mutation(s) at GyrA | Mutation(s) at ParC |
| --- | --- | --- | --- | --- | --- |
| *K. pneumoniae* ATCC 10031 wt | E2 | Ciprofloxacin | 116 | - | - |
| *K. pneumoniae* ATCC 10031 wt | E4 | Ciprofloxacin | 116 | D87Y | - |
| *K. pneumoniae* ATCC 10031 wt | E6 | Ciprofloxacin | 116 | D87Y | - |
| *K. pneumoniae* ATCC 10031 wt | F3 | Ciprofloxacin | 116 | - | - |
| *K. pneumoniae* ATCC 10031 wt | F5 | Ciprofloxacin | 80 | - | - |
| *K. pneumoniae* ATCC 10031 wt | F7 | Ciprofloxacin | 100 | - | - |
| *E. coli* K-12 MG1655 wt | E8 | Ciprofloxacin | 116 | **D82G**, S83D | - |
| *E. coli* K-12 MG1655 wt | E10 | Ciprofloxacin | 116 | D87N | - |
| *E. coli* K-12 MG1655 wt | F9 | Ciprofloxacin | 100 | S83L, D87G | E84K, N105S |
| *E. coli* K-12 MG1655 wt | F11 | Ciprofloxacin | 116 | S83L, D87G | E84K, N105S |
| *E. coli* K-12 MG1655 wt | G2 | Ciprofloxacin | 68 | S83L, D87G | E84K |
| *E. coli* K-12 MG1655 wt | H3 | Ciprofloxacin | 84 | S83L, D87G | E84K |
| *E. coli* K-12 MG1655 $\Delta mutS$ | G4 | Ciprofloxacin | 72 | S83L, D87A | S80I |
| *E. coli* K-12 MG1655 $\Delta mutS$ | G6 | Ciprofloxacin | 64 | S83L, D87G | E84K |
| *E. coli* K-12 MG1655 $\Delta mutS$ | G8 | Ciprofloxacin | 76 | S83L, A119E | G78D |
| *E. coli* K-12 MG1655 $\Delta mutS$ | H5 | Ciprofloxacin | 72 | S83L, D87G | E84G |
| *E. coli* K-12 MG1655 $\Delta mutS$ | H7 | Ciprofloxacin | 72 | S83L, D87G | S80I, E84G |
| *E. coli* K-12 MG1655 $\Delta mutS$ | H9 | Ciprofloxacin | 72 | S83L, D87G | **D79N**, S80R |

**Supplementary Table S3.** Sequence analysis of DIvERGE-generated mutants of *Klebsiella pneumoniae* ATCC 10031 conferring resistance to gepotidacin.

| Library | Clone ID | Mutation at GyrA | Mutation at ParC |
| --- | --- | --- | --- |
| **DivERGE mutagenesis of *K. pneumoniae* GyrA and GyrB** | #1 | D82N | D79N |
|  | #2 | D82N | D79N |
|  | #3 | D82N | D79N |
|  | #4 | D82N | D79N |
|  | #5 | D82N | D79N |
|  | #6 | D82N | D79N |
|  | #7 | D82N | D79N |
|  | #8 | D82N | D79N |
|  | #9 | D82N | D79N |
|  | #10 | D82N | D79N |
| **Saturation mutagenesis of *K. pneumoniae* GyrA Asp82 and ParC Asp79** | #1 | D82N | D79N |
|  | #2 | D82N | D79N |
|  | #3 | D82N | D79N |
|  | #4 | D82N | D79N |
|  | #5 | D82N | D79N |
|  | #6 | D82N | D79N |
|  | #7 | D82N | D79N |
|  | #8 | D82N | D79N |
|  | #9 | D82N | D79N |
|  | #10 | D82N | D79N |

**Supplementary Figure S4.**
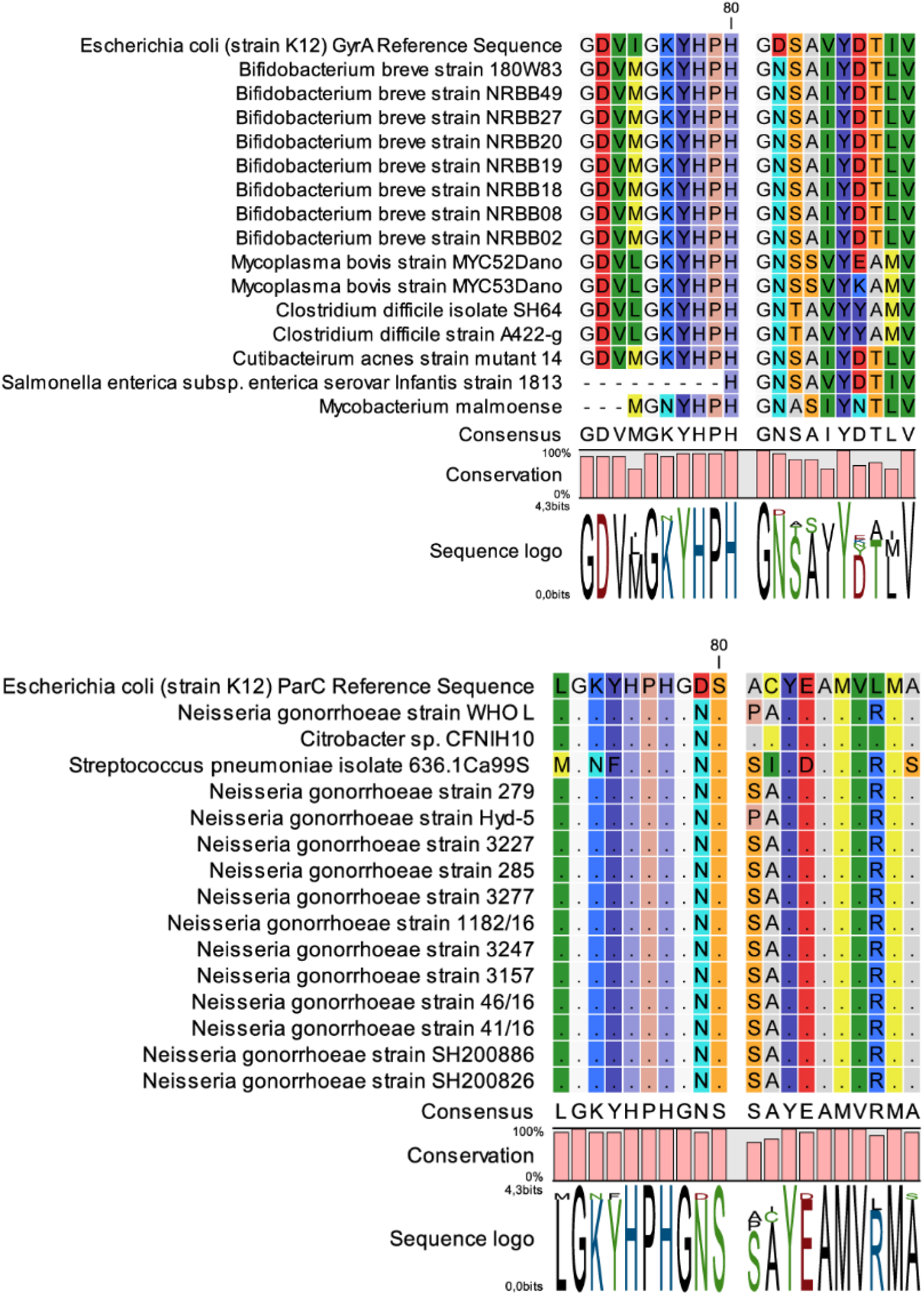
tBLASTn analysis identified gepotidacin resistance-associated genotypes in the nucleotide archive of the National Center for Biotechnology Information (NCBI). See Supplementary File 1 for taxon identifiers, GeneBank IDs, and analyzed protein sequences.

## Materials and Methods

### Media and antibiotics

For general growth of bacteria and electrocompetent cell preparation, Lysogeny-Broth-Lennox (LB), medium was used (10 g tryptone, 5 g yeast extract, 5 g sodium chloride per 1 l of water). For cell recovery after electroporation, we applied Terrific Broth (TB) (24 g yeast extract, 12 g tryptone, 9.4 g K_2_HPO_4_, and 2 g KH_2_PO_4_ per 1 l of water). For antimicrobial susceptibility tests and the selection of resistant variants, cation-adjusted Mueller Hinton II Broth (MHBII) was used. To prepare MHBII broth, 22 g of MHBII powder (Becton Dickinson and Co.) was dissolved in 1 l of water (3 g beef extract, 17.5 g acid hydrolysate of casein and 1.5 g starch). MHBII agar was prepared by the addition of 14 g Bacto agar to 1 l bacterial MHBII broth. Bacterial media were sterilized by autoclaving 15 min at 121 °C. To distinguish lacZ knockout strains and wild-type strains, sterile solutions of X-gal (5-bromo-4-chloro-3-indolyl-β-D-galactopyranoside) and IPTG (isopropyl β-D-1-thiogalactopyranoside) were added to the MHBII agar medium after autoclaving. Final X-gal and IPTG concentration in the medium were 40 g/l and 0.2 μM, respectively. Antibiotics and chemicals were ordered from Sigma-Aldrich (ampicillin, kanamycin, chloramphenicol; X-gal, IPTG), from Fluka Analytical (ciprofloxacin), and MedChemExpress (gepotidacin).

### Oligonucleotides

Sequences of all synthetic DNA oligonucleotides (oligos) are listed in Supplementary file 1. Oligonucleotides were synthesized at the Nucleic Acid Synthesis Laboratory of the Biological Research Centre of the Hungarian Academy of Sciences. Soft-randomized DIvERGE oligos were manufactured according to a previously described soft-randomization protocol^22^. PCR primers were purified with standard desalting, while mutagenic oligos were purified with high-performance liquid chromatography (HPLC). Lyophilized oligos were dissolved in 1× Tris-EDTA (TE) buffer, pH 8 (Integrated DNA Technologies) at a final concentration of 100 μM. DIvERGE oligos were dissolved in 0.5× TE buffer at a final concentration of 500 μM. Dissolved oligos were stored at −20 °C.

### Computational docking of gepotidacin

We docked gepotidacin (ChemSpider ID: 34982930) into the DNA gyrase and topoisomerase IV homology models using Glide from the Schrödinger package^70^. The protonation state of gepotidacin was selected as the main species obtained from pKa calculation using the calculator plugin of Marvin program^71^. For Glide docking we used OPLS3 force field^72^ and the standard precision procedure with flexible ligand geometries and enhanced conformational sampling. Based on the high similarity of the mode-of-action of GSK945237 and gepotidacin^61^, we picked the docking pose of gepotidacin that was most similar to the binding pose of GSK945237 in 5NPP structure. This procedure was repeated for the other docking-pose of GSK945237, as well. Finally, the binding site (defined as residues closer than 5 Å to the ligand) geometry together with the docked ligand was optimized. We followed the same steps in the case of both homology models.

### Molecular dynamics simulations of gepotidacin bound DNA-gyrase and DNA-topoisomerase IV complexes

To perform molecular dynamics-based binding mode analysis for gepotidacin at both of its targets, we first parametrized the complex of formed by gepotidacin-DNA- and its corresponding target (DNA gyrase and topoisomerase IV, respectively). For parametrization, we used OPLS3 force field^72^. Next, this system was solvated using the SPC (simple point charge) explicit water model^73^ within an orthorhombic box 10 Å apart. On the course of simulations, NaCl in 0.15 M final concentration was used to mimic the physiological conditions. 100 ns molecular dynamics calculations were carried out on these systems at constant volume and temperature (NVT) using the Desmond molecular dynamics code^79^. Snapshots of these simulations were used as a conformational ensemble in mutation-induced binding free energy change calculations.

### Estimation of mutation-induced binding free energy changes

Asparagine-mutation induced changes of the binding affinity (ΔΔG_B_) of gepotidacin at GyrA D82, and ParC D79 positions were calculated as the difference of their binding free energy in the mutated and wild-type enzyme (ΔΔG_B_= ΔG_B_^MUT^ -ΔG_B_^WT^). This affinity change was estimated by using Residue Scanning module of Bioluminate package^75^ and was based on molecular mechanics-generalized Born surface area (MM-GBSA) approximation^21^. Specifically, we estimated the binding free energy of a ligand (Δ*G*)by using the single trajectory approach^76^ in which a single molecular dynamics simulation of the protein-ligand complex is completed first and sample geometries are collected from the trajectory to represent the possible binding conformations. Δ*G* is calculated for every single geometry as

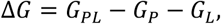

where *G*_*PL*_, *G*_*P*_ and *G*_*L*_ are the energy of the protein ligand complex, protein and ligand, respectively. These energy values are calculated as

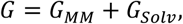

where *G*_*MM*_ is the calculated molecular mechanics energy for the force field applied, and *G*_*Solv*_ is the solvation energy in the generalized Born approximation. Different variations of this method are widely used in large scale drug design studies^81–83^ as it accurately predicts mutational effects in drug-protein interaction analyses with a manageable computational demand (see *e.g*. references^80,81^). In our study, MM-GBSA calculations were carried out with the *thermal_mmgbsa.py* script Schrödinger package^75^ from 100, evenly spaced structures from the second 50 ns of our previously performed molecular-dynamics simulation between the wild-type enzyme (DNA gyrase and topoisomerase IV) and gepotidacin.

### Plasmid construction

To perform ssDNA recombineering and DIvERGE in *K. pneumoniae* ATCC 10031, we generated a novel version of pORTMAGE, termed pORTMAGE311B. This plasmid possesses a tightly regulated, XylS-Pm regulator/promoter-based λ Beta and *E. coli* MutL E32K co-expression system. On, pORTMAGE the expression of MutL E32K of *E. coli* abolishes methyl-directed mismatch repair (MMR) during the time-period of oligonucleotide incorporation into the bacterial genome^60^. pORTMAGE311B ensured functionality at 37 °C and the rapid inducibility of λ Beta and E. coli MutL E32K by the addition of 1 mM m-Toluic-acid as an inducer. The temporal co-expression of λ Beta and MutL E32K also limits off-target mutagenic effects on the course of genome editing. Construction of pORTMAGE311B was performed by introducing the coding sequence of the dominant *E. coli mutL*(E32K) allele to the pSEVA258*beta* plasmid^82^. To this aim, *mutL*(E32K) was PCR amplified from pORTMAGE2^60^ using primers RBE32KF and XE32KR. The PCR product was then purified and digested with BamHI and XbaI. Next, the resulted fragment was cloned downstream of *beta* within pSEVA258*beta*. Finally, the successful assembly of the plasmid was verified by PCR and confirmed by capillary sequencing. pORTMAGE311B was also deposited to Addgene (Addgene Number 120418).

### DIvERGE in *Klebsiella pneumoniae*

DIvERGE in *Klebsiella pneumoniae* ATCC 10031 was carried out according to a previously described DIvERGE workflow^22^, with minor modifications. Briefly, *K. pneuomoniae* cells carrying pORTMAGE311B plasmid were inoculated into 2 ml LB medium + 50 μg/ml kanamycin and were grown at 37 °C, 250 rpm overnight. From this starter culture, 500 μl stationary phase culture was inoculated into 50 ml LB medium + 50 μg/ml kanamycin, and grown at 37 °C under continuous shaking at 250 rpm. Induction was initiated at OD_600_ = 0.4 by adding 50 μl 1 M m-Toluic-acid (Sigma-Aldrich, dissolved in 96% ethyl-alcohol). Induction was performed for 30 minutes at 37 °C. After induction, cells were cooled on ice for 15 minutes. Next, cells were washed 3-times with 50 ml sterile, ice-cold, ultra-pure distilled water. Finally, the cell pellet was resuspended in 800 μl sterile ultra-pure distilled water and kept on ice until electroporation.

To perform DIvERGE, and separately, the saturation mutagenesis of GyrA D82 and ParC D79, the corresponding *gyrA,* and *parC-*targeting oligos were equimolarly mixed at a final concentration of 500 μM and 2 μl of this oligo-mixture was added to 40 μl electrocompetent cell in 10 parallels. Following the electroporation of each sample, the 10 parallel samples were pooled together, and immediately after electroporation cells were suspended in 50 ml TB to allow for cell recovery. After a 1-hour recovery period at 37 °C, 250 rpm, an additional 50 ml LB medium along with 50 μg/ml (final cc.) kanamycin was added. Next, cells were grown at 37 °C, 250 rpm for 24 hours. To select gepotidacin resistant clones from this DIvERGE library, 500 μl cell library was spread to 6 μg/ml gepotidacin-containing MHBII agar plates and plates were incubated at 37 °C for 48 hours. Finally, 10 randomly selected colonies were analyzed by capillary sequencing (oligos: KPGA1F, KPGA1R and KPPC1F, KPPC1R) from both antibiotic-selected cell libraries.

### Engineering gepotidacin resistance-associated mutations in *Escherichia coli*

To assess the phenotype of gepotidacin resistance-associated mutations, we individually reconstructed both GyrA D82N and ParC D79N mutations in *E. coli* K-12 MG1655. We utilized a previously described CRISPR-MAGE protocol to integrate mutations and counter select against the wild-type genotype^83,84^. Briefly, cell-containing pORTMAGE2 (Addgene #72677) were transformed with pCas9 (Addgene #42876) and were grown in LB + 100 μg/ml ampicillin + 20 μg/ml chloramphenicol broth at 30 °C. Next, pCRISPR plasmids (Addgene #42875) were constructed with crRNA sequences targeting the wild-type *gyrA* and *parC* loci in the vicinity of D82 and D79 according to a previously described protocol (oligos: ECGACRF1, ECGACRR1, and ECPCCRF1, ECPCCRR1, respectively). Following plasmid-construction, correct clones were identified by capillary sequencing. To integrate GyrA D82N and ParC D79N mutations into the chromosome of *E. coli*, induced pORTMAGE2 and pCas9-carrying cells were prepared and made electrocompetent. Next, simultaneously, 100 ng of the corresponding pCRISPR plasmid and 200 pmole of oligonucleotide (MGYRA82N or MPARC79N) per 40 μl electrocompetent cells was electroporated. Cell were allowed to recover in 1 ml TB media and grown in 5 ml LB + 100 μg/ml ampicillin + 20 μg/ml chloramphenicol broth at 30 °C overnight and then spread to LB + 50 μg/ml kanamycin + 20 μg/ml chloramphenicol agar plates. Finally, the presence of the correct mutations was identified by capillary sequencing of the oligo-targets (oligos: 25922Ga1, 25922Ga3, and 25922Pc1, 25922Pc3).

### Engineering isogenic *Klebsiella* and *Salmonella* strains carrying gepotidacin and fluoroquinolone resistance-associated mutations

To investigate mutational effects on antibiotic susceptibility and growth phenotypes, gepotidacin and ciprofloxacin resistance-linked mutations and mutation combinations were reconstructed in *K. pneumoniae* ATCC 10031 and in *Salmonella enterica ssp. enterica* serovar *Typhimurium*. To this aim, specific point mutations (see Supplementary file 1) were inserted into the chromosome of *K. pneumoniae* by pORTMAGE-recombineering^22,60^. Briefly, pORTMAGE-recombineering was performed using pORTMAGE311B (Addgene #120418). 1 μl of 100 μM of the corresponding oligos or oligo-mixtures were used for ssDNA-recombineering in appropriate combinations to create ciprofloxacin and gepotidacin resistance-conferring mutations. Following recombineering, cells were allowed to grow overnight in TB medium at 37°C, 250 rpm. Next, variants carrying ciprofloxacin resistance-conferring mutations were selected on LB + 100 ng/ml ciprofloxacin agar plates, while mutants carrying the GyrA D82N; ParC D79N mutation-combination were selected on MHBII + 6 μg/ml gepotidacin agar plates. Cells carrying individual gepotidacin resistance-associated mutations were plated onto LB + 50 μg/ml kanamycin agar plates. To obtain individual colonies, cultures from each recombineering populations were diluted, and appropriate dilutions were spread to agar plates. Plates were incubated at 37 °C, and individual colonies were genotyped by allele-specific PCR (using oligos KPA82ASF, KPA82ASR, and KPC79ASF, KPC79ASR). Finally, positive clones were confirmed by capillary sequencing of the oligo-target region. The reconstruction of the clinically prevalent GyrA D82 mutation in combination with a ParC D79N mutation in *Salmonella enterica ssp. enterica* serovar *Typhimurium* LT2 were conducted similarly to the reconstruction of specific mutations in *K. pneumoniae*. Briefly, pORTMAGE311B-containing, induced *S. enterica* cells were made electrocompetent and transformed with 1 μl of SE_GA_D82N and SE_PC_D79N oligos that integrated the corresponding GyrA D82N and ParC D79N mutations into the bacterial chromosome. Next, mutants were selected on MHBII + 6 μg/ml gepotidacin agar plates at 37 °C. Finally, the presence of the two mutations was identified and validated by capillary sequencing using SEGAF1, SEGAR1, and SEPCF1, SEPCR1 oligos.

### Adaptive laboratory evolution of ciprofloxacin resistance

Adaptive laboratory evolution experiments followed an established protocol for automated laboratory evolution^85^ and aimed to maximize the drug-resistance increment during a fixed time period. At each transfer step, 10^7^ bacterial cells were transferred to a new culture and adaptation were performed by passaging 6 independent populations of *Klebsiella pneumoniae* ATCC 10031 wild-type, *Escherichia coli* K-12 MG1655 wild-type, and *Escherichia coli* K-12 MG1655 Δ*mutS* in the presence of increasing ciprofloxacin concentrations. Experiments was conducted in 96-well plates, in MHBII medium, by utilizing a checker board layout to minimize and to monitor cross-contamination. These 96-well deep-well plates (0.5 ml, polypropylene, V-bottom) were covered with sandwich covers (Enzyscreen BV) to ensure an optimal oxygen exchange rate and limit evaporation and were shaken at 150 rpm at 37 °C. Twenty μl of each evolving culture was parallelly transferred into four independent wells containing 350 μl fresh medium and an increasing concentration of gepotidacin and ciprofloxacin (i.e., 0.5×, 1×, 1.5×, and 2.5× the concentration of the previous concentration step). Following cell transfer, each culture was allowed to grow for 48 h. At each transfer, cell growth was monitored by measuring the optical density at 600 nm (OD_600_) (Biotek Synergy 2). Only populations with (i) vigorous growth (i.e., OD_600_ > 0.25) and (ii) the highest drug concentration were selected for further evolution. Accordingly, only one of the four populations was retained for each independently evolving strain. This protocol was designed to avoid population extinction and to ensure that populations with the highest level of resistance were propagated further during evolution. Samples from each transfer were frozen in 15% dimethyl-sulfoxide (DMSO) at -80 °C for further analysis. Adaptation of an individual population was terminated when the antibiotic concentration in the given well would have exceeded 200 μg/ml after the transfer. Cells from these, highly-drug resistant populations were frozen after the addition of 15% dimethyl-sulfoxide (DMSO) and were kept at −80 °C. Following adaptation, cells from each final population were spread onto MHBII agar plates and individual colonies were isolated. Next, 3 colonies from each adapted line were subjected to capillary sequencing at *gyrA* and *parC* to assess their genotype. Target regions were amplified in *K. pneumoniae* by using KPGA1F, KPGA1R, and KPPC1F, KPPC1R oligos, while for *E. coli* 25922Ga1, 25922Ga3, and 25922Pc1, 25922Pc3 were applied. Capillary sequencing indicated that all colonies within the given adapted population were isogenic in all cases.

### *In vitro* growth-rate measurements

We measured the growth phenotype of bacterial variants by assessing their growth at 37°C in MHBII medium. To measure growth, we inoculated 5×10^4^ cells from early stationary phase cultures (prepared in MHBII medium) into 100 μl of MHBII medium in a 96 well microtiter plate and monitored growth for 24 h. Bacterial growth was measure as the 600 nm optical density (OD_600_) of cultures at a given timepoint. OD_600_ measurements were carried out every 5 minutes using BioTek Synergy 2 microplate reader while bacterial cultures were grown at 37°C under continuous, variable intensity shaking. Each bacterial variant and their corresponding wild-types were measured in two consecutive experiments, with 12 replicates in each subsequent tests (24 biological replicates in total). Finally, growth rates were calculated from the obtained growth curves according to a previously described procedure^86,87^.

### Competition-based fitness measurements

Competition assay-based fitness measurements were carried out by competing mutant strains against their corresponding wild-type strain carrying a *lacZ* knockout mutation. The *lacZ* knockout strain was constructed by integrating a premature stop codon in place the 23^rd^ amino acid of LacZ with a previously reported pORTMAGE protocol^60^. Briefly, heat-induced pORTMAGE3 (Addgene #72678) containing *K. pneuomoniae* ATCC 10031 cells were electroporated with KpLacZW23* oligonucleotide (2.5 nM final concentration). Following oligo-integration into the bacterial chromosome, *lacZ*(-) variants were identified by plating cells out to X-gal + IPTG-containing MHBII agar plates at appropriate dilution to form single colonies, where knockout mutants formed white colonies and could be easily distinguished from the dark blue colonies of the wild-type containing a functional β-galactosidase gene. Each competition experiment started by inoculating early stationary phase cultures in a 99:1, as mutant to wild-type ratio, into 10 ml MHBII medium at 1:1000-fold dilution and incubating each culture at 37 °C for 24 h under a constant agitation of 250 rpm. These cultures were then serially transferred into 10 ml fresh MHBII medium in a 1:1000 dilution in every 24 hours for 3 subsequent transfers. To analyze the composition of each population, cultures were plated onto X-gal + IPTG supplemented MHBII agar plates (in 145×20 mm Petri dishes, Greiner Bio-One Ltd) at appropriate dilutions to obtain approximately 1000 colonies per each plate. Plates were then incubated at 37 °C overnight. The ratio of the mutant (*lacZ* proficient, blue) and the wild-type (*lacZ* deficient, white) at each given time point was determined from the number of blue and white colonies on X-Gal + IPTG agar plates. Importantly, prior experiments showed that *lacZ* deficient K. pneumoniae cells suffer no competitive disadvantage compere to their corresponding wild-type strain. Competition experiments were performed in five replicates. Finally, selection coefficients were estimated as the slope of the change in ratio, as defined in the following equation:

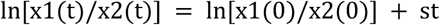

where s is the selection coefficient, t is time or number of generations, x1 and x2 are the ratio of the two strains at a given time point^88^.

### Prevalence of gepotidacin resistance-associated mutations in sequence databases

In order to investigate if genotypes GyrA D82 and ParC D79N exist in bacteria, we downloaded all DNA sequences from NCBI nucleotide database (ftp://ftp.ncbi.nlm.nih.gov/blast/db/nt.??.tar.gz) using wget 1.17.1 on 9 Oct 2018. We performed the BLAST search with tBLASTn 2.231^89^ (build Jan 7 2016) on the downloaded full nucleotide database as subject and GyrA and ParC as query. Protein sequences of GyrA (P0AES4) and ParC (P0AFI2) were downloaded from UniProt. To perform BLAST, the database was fragmented and the BLAST search was carried out on these smaller datasets (with argument max_target_seqs = 20000). Finally, separately obtained results were merged. Finally, for each sequence the taxonomy of the source bacteria was assigned with blastdbcmd (version 2.2.31).

### Mouse thigh infection models

Drug-resistant *K. pneumoniae* is frequently responsible for wound and systemic infections, therefore, we benchmarked *in vivo* effects of gepotidacin resistance in a murine thigh wound infection model^31,32^. Specifically, we examined wound colonization for representative gepotidacin and ciprofloxacin-resistant mutants of *K. pneumoniae* ATCC 10031 and compared to that of the wild-type. Murine thigh infections were performed according to a previously established protocol^32,90^. Bacterial inoculums were prepared by inoculating single bacterial colonies into 5 ml of MHBII broth and were incubated at 37 °C for 16 hours under constant agitation (250 rpm). Next, ½ volume of 50 volume/volume% glycerol:water mixture was added to each culture and 550 μl of these cell suspensions were frozen at −80 °C. Before inoculation, cell suspensions were allowed to thaw at room temperature for 15 minutes. As test animals, groups of 5 female specific-pathogen-free ICR (CD-1) mice weighing 22 ± 2 g were used. Animals were immunosuppressed by two intraperitoneal injections of cyclophosphamide, the first at 150 mg/kg 4 days before infection (day –4) and the second at 100 mg/kg 1 day before infection (day –1). On day 0, animals were inoculated intramuscularly into the right thigh with 10^6^ CFU/mouse of the corresponding pathogenic mutant (0.1 ml culture/thigh). After 26 hours inoculation, animals were humanely euthanized with CO_2_ asphyxiation and then the muscle of the right thigh was harvested from each test animal. The removed muscle was homogenized in 3 ml of phosphate-buffered saline, pH 7.4, with a polytron homogenizer. Finally, 0.1 mls of these homogenates were used for serial 10-fold dilutions and plated onto LB agar for colony count (CFU/g) determination.

All aspects of this work including housing, experimentation, and animal disposal were performed in general accordance with the Guide for the Care and Use of laboratory animals (National Academy Press, Washington, DC, 2011). All experiments were performed under ABSL2 conditions in the AAALAC(American Association for Accreditation of Laboratory Animal Care)-accredited vivarium of Eurofins Pharmacology Discovery Services Taiwan, Ltd. with the oversight of veterinarians to assure compliance with IACUC regulations and the humane treatment of laboratory animals.

### Minimum Inhibitory Concentration measurements

Minimum Inhibitory Concentrations (MICs) were determined using a standard serial broth microdilution technique according to the EUCAST guidelines^23^ (ISO 20776-1:2006, Part 1). Briefly, bacterial strains were inoculated from frozen cultures onto MHBII agar plates and were grown overnight at 37 °C. Next, independent colonies from each strain were inoculated into 1 ml MHBII medium and were propagated at 37 °C, 250 rpm overnight. To perform MIC assays, 12-step serial dilutions using 2-fold dilution-steps of the given antibiotic were generated in 96-well microtiter plates (Sarstedt 96-well microtest plate with lid, flat base). Antibiotics were diluted in 100 μl of MHBII medium. Following dilutions, each well was seeded with an inoculum of 5×10^4^ bacterial cells. Each measurement were performed at least in 3 parallel replicates. Also, to avoid possible edge effects at the edge of the microwell plate, side-rows (A and H) contained only medium without bacterial cells. Following inoculations, plates were covered with lids and wrapped in polyethylene plastic bags to minimize evaporation but allow for aerobic O_2_ transfer. Plates were incubated at 37 °C under continuous shaking at 150 rpm for 18 hours in an INFORS HT shaker. After incubation, OD_600_ of each well on the microwell plate was measured using a Biotek Synergy 2 microplate reader. MIC was defined as the antibiotic concentration which inhibited the growth of the bacterial culture, i.e., the drug concentration where the average OD_600_ increment of the three replicates was below 0.05.

## References

1. Silver, L. L. Multi-targeting by monotherapeutic antibacterials. Nat Rev Drug Discov 6, 41–55 (2007).

2. Spellberg, B., Powers, J. H., Brass, E. P., Miller, L. G. & Edwards, J. E. Trends in Antimicrobial Drug Development: Implications for the Future. Clin Infect Dis. 38, 1279–1286 (2004).

3. Jackson, N., Czaplewski, L. & Piddock, L. J. V. Discovery and development of new antibacterial drugs: learning from experience? J Antimicrob Chemother doi:10.1093/jac/dky019

4. Silver, L. L. Challenges of Antibacterial Discovery. Clin. Microbiol. Rev. 24, 71–109 (2011).

5. O’Dwyer, K. et al. Bacterial Resistance to Leucyl-tRNA Synthetase Inhibitor GSK2251052 Develops during Treatment of Complicated Urinary Tract Infections. Antimicrob. Agents Chemother. 59, 289–298 (2015).

6. Suzuki, S., Horinouchi, T. & Furusawa, C. Suppression of antibiotic resistance acquisition by combined use of antibiotics. Journal of Bioscience and Bioengineering (2015). doi:10.1016/j.jbiosc.2015.02.003

7. Munck, C., Gumpert, H. K., Wallin, A. I. N., Wang, H. H. & Sommer, M. O. A. Prediction of resistance development against drug combinations by collateral responses to component drugs. Sci Transl Med 6, 262ra156–262ra156 (2014).

8. Klahn, P. & Brönstrup, M. Bifunctional antimicrobial conjugates and hybrid antimicrobials. Nat. Prod. Rep. (2017). doi:10.1039/C7NP00006E

9. Farrell, D. J., Sader, H. S., Rhomberg, P. R., Scangarella-Oman, N. E. & Flamm, R. K. In Vitro Activity of Gepotidacin (GSK2140944) against Neisseria gonorrhoeae. Antimicrob. Agents Chemother. 61, e02047–16 (2017).

10. Dougherty, T. J. et al. NBTI 5463 Is a Novel Bacterial Type II Topoisomerase Inhibitor with Activity against Gram-Negative Bacteria and In Vivo Efficacy. Antimicrobial Agents and Chemotherapy 58, 2657–2664 (2014).

11. Tari, L. W. et al. Tricyclic GyrB/ParE (TriBE) Inhibitors: A New Class of Broad-Spectrum Dual-Targeting Antibacterial Agents. PLOS ONE 8, e84409 (2013).

12. Wang, K. K. et al. A Hybrid Drug Limits Resistance by Evading the Action of the Multiple Antibiotic Resistance Pathway. Mol Biol Evol 33, 492–500 (2016).

13. Pokrovskaya, V., Belakhov, V., Hainrichson, M., Yaron, S. & Baasov, T. Design, Synthesis, and Evaluation of Novel Fluoroquinolone−Aminoglycoside Hybrid Antibiotics. J. Med. Chem. 52, 2243–2254 (2009).

14. Nayar, A. S. et al. Target-Based Resistance in Pseudomonas aeruginosa and Escherichia coli to NBTI 5463, a Novel Bacterial Type II Topoisomerase Inhibitor. Antimicrobial Agents and Chemotherapy 59, 331–337 (2015).

15. Taylor, S. N. et al. Gepotidacin for the Treatment of Uncomplicated Urogenital Gonorrhea: A Phase 2, Randomized, Dose-Ranging, Single-Oral Dose Evaluation. Clin Infect Dis doi:10.1093/cid/ciy145

16. O’Riordan, W. et al. The Efficacy, Safety, and Tolerability of Gepotidacin (GSK2140944) in the Treatment of Patients with Suspected or Confirmed Gram-Positive Acute Bacterial Skin and Skin Structure Infections. Antimicrobial Agents and Chemotherapy AAC.02095-16 (2017). doi:10.1128/AAC.02095-16

17. Biedenbach, D. J. et al. In Vitro Activity of Gepotidacin, a Novel Triazaacenaphthylene Bacterial Topoisomerase Inhibitor, against a Broad Spectrum of Bacterial Pathogens. Antimicrob. Agents Chemother. 60, 1918–1923 (2016).

18. Flamm, R. K., Farrell, D. J., Rhomberg, P. R., Scangarella-Oman, N. E. & Sader, H. S. Gepotidacin (GSK2140944) In Vitro Activity against Gram-positive and Gram-negative Bacteria (MBC/MIC, Kill Kinetics, Checkerboard, PAE/SME tests). Antimicrob. Agents Chemother. AAC.00468-17 (2017). doi:10.1128/AAC.00468-17

19. Nyerges, Á. et al. Directed evolution of multiple genomic loci allows the prediction of antibiotic resistance. PNAS 201801646 (2018). doi:10.1073/pnas.1801646115

20. Tillotson, G. A crucial list of pathogens. The Lancet Infectious Diseases 0, (2017).

21. Kollman, P. A. et al. Calculating Structures and Free Energies of Complex Molecules:?Combining Molecular Mechanics and Continuum Models. Acc. Chem. Res. 33, 889–897 (2000).

22. Nyerges, Á. et al. Directed evolution of multiple genomic loci allows the prediction of antibiotic resistance. PNAS 115, E5726–E5735 (2018).

23. ISO 20776-1:2006 -Clinical laboratory testing and in vitro diagnostic test systems - Susceptibility testing of infectious agents and evaluation of performance of antimicrobial susceptibility test devices - Part 1: Reference method for testing the in vitro activity of antimicrobial agents against rapidly growing aerobic bacteria involved in infectious diseases. ISO Available at: http://www.iso.org/iso/catalogue_detail.htm?csnumber=41630. (Accessed: 5th February 2017)

24. Hughes, D. & Andersson, D. I. Evolutionary consequences of drug resistance: shared principles across diverse targets and organisms. Nat Rev Genet 16, 459–471 (2015).

25. Martínez, J. L., Baquero, F. & Andersson, D. I. Beyond serial passages: new methods for predicting the emergence of resistance to novel antibiotics. Current Opinion in Pharmacology 11, 439–445 (2011).

26. Sommer, M. O. A., Munck, C., Toft-Kehler, R. V. & Andersson, D. I. Prediction of antibiotic resistance: time for a new preclinical paradigm? Nat Rev Micro advance online publication, (2017).

27. Andersson, D. I. Improving predictions of the risk of resistance development against new and old antibiotics. Clinical Microbiology and Infection 21, 894–898 (2015).

28. Krapp, F., Ozer, E. A., Qi, C. & Hauser, A. R. Case Report of an Extensively Drug-Resistant Klebsiella pneumoniae Infection With Genomic Characterization of the Strain and Review of Similar Cases in the United States. Open Forum Infect Dis 5, (2018).

29. Weigel, L. M., Steward, C. D. & Tenover, F. C. gyrA Mutations Associated with Fluoroquinolone Resistance in Eight Species ofEnterobacteriaceae. Antimicrob. Agents Chemother. 42, 2661–2667 (1998).

30. Chen, F.-J., Lauderdale, T.-L., Ho, M. & Lo, H.-J. The Roles of Mutations in gyrA, parC, and ompK35 in Fluoroquinolone Resistance in Klebsiella pneumoniae. Microbial Drug Resistance 9, 265–271 (2003).

31. Velkov, T., Bergen, P., Lora-Tamayo, J., Landersdorfer, C. B. & Li, J. PK/PD models in antibacterial development. Curr Opin Microbiol 16, (2013).

32. Tan, C. M. et al. Restoring Methicillin-Resistant Staphylococcus aureus Susceptibility to β-Lactam Antibiotics. Science Translational Medicine 4, 126ra35–126ra35 (2012).

33. Hooper, D. C. & Jacoby, G. A. Topoisomerase Inhibitors: Fluoroquinolone Mechanisms of Action and Resistance. Cold Spring Harb Perspect Med 6, a025320 (2016).

34. Gibson, E. G., Ashley, R. E., Kerns, R. J. & Osheroff, N. Bacterial Type II Topoisomerases and Target-Mediated Drug Resistance. in Antimicrobial Resistance in the 21st Century (eds. Fong, I. W., Shlaes, D. & Drlica, K.) 507–529 (Springer International Publishing, 2018). doi:10.1007/978-3-319-78538-7_16

35. Wohlkonig, A. et al. Structural basis of quinolone inhibition of type IIA topoisomerases and target-mediated resistance. Nat Struct Mol Biol 17, 1152–1153 (2010).

36. Collin, F., Karkare, S. & Maxwell, A. Exploiting bacterial DNA gyrase as a drug target: current state and perspectives. Appl Microbiol Biotechnol 92, 479–497 (2011).

37. Correia, S., Poeta, P., Hébraud, M., Capelo, J. L. & Igrejas, G. Mechanisms of quinolone action and resistance: where do we stand? Journal of Medical Microbiology 66, 551–559 (2017).

38. Randall, L. P., Coldham, N. G. & Woodward, M. J. Detection of mutations in Salmonella entericagyrA, gyrB, parC and parE genes by denaturing high performance liquid chromatography (DHPLC) using standard HPLC instrumentation. J Antimicrob Chemother 56, 619–623 (2005).

39. Eaves, D. J. et al. Prevalence of Mutations within the Quinolone Resistance-Determining Region of gyrA, gyrB, parC, and parE and Association with Antibiotic Resistance in Quinolone-Resistant Salmonella enterica. Antimicrobial Agents and Chemotherapy 48, 4012–4015 (2004).

40. Thong, K. L., Ngoi, S. T., Chai, L. C. & Teh, C. S. J. Quinolone Resistance Mechanisms Among Salmonella enterica in Malaysia. Microbial Drug Resistance 22, 259–272 (2015).

41. García-Fernández, A. et al. Emergence of Ciprofloxacin-Resistant Salmonella enterica Serovar Typhi in Italy. PLOS ONE 10, e0132065 (2015).

42. Su, X. & Lind, I. Molecular Basis of High-Level Ciprofloxacin Resistance in Neisseria gonorrhoeae Strains Isolated in Denmark from 1995 to 1998. Antimicrob. Agents Chemother. 45, 117–123 (2001).

43. Jacobsson, S., Golparian, D., Scangarella-Oman, N. & Unemo, M. In vitro activity of the novel triazaacenaphthylene gepotidacin (GSK2140944) against MDR Neisseria gonorrhoeae. J Antimicrob Chemother doi:10.1093/jac/dky162

44. Hamasuna, R. et al. Mutations in ParC and GyrA of moxifloxacin-resistant and susceptible Mycoplasma genitalium strains. PLOS ONE 13, e0198355 (2018).

45. Friedman, S. M., Lu, T. & Drlica, K. Mutation in the DNA Gyrase A Gene of Escherichia coli That Expands the Quinolone Resistance-Determining Region. Antimicrob Agents Chemother 45, 2378–2380 (2001).

46. Chu, Y.-W., Houang, E. T. S. & Cheng, A. F. B. Novel Combination of Mutations in the DNA Gyrase and Topoisomerase IV Genes in Laboratory-Grown Fluoroquinolone-Resistant Shigella flexneri Mutants. Antimicrobial Agents and Chemotherapy 42, 3051–3052 (1998).

47. Minnick, M. F., Wilson, Z. R., Smitherman, L. S. & Samuels, D. S. gyrA Mutations in Ciprofloxacin-Resistant Bartonella bacilliformis Strains Obtained In Vitro. Antimicrob. Agents Chemother. 47, 383–386 (2003).

48. Bachoual, R., Dubreuil, L., Soussy, C.-J. & Tankovic, J. Roles of gyrA Mutations in Resistance of Clinical Isolates and In Vitro Mutants of Bacteroides fragilis to the New Fluoroquinolone Trovafloxacin. Antimicrob Agents Chemother 44, 1842–1845 (2000).

49. Scangarella-Oman, N. E. et al. Microbiological Analysis From a Phase 2 Randomized Study in Adults Evaluating Single Oral Doses of Gepotidacin in the Treatment of Uncomplicated Urogenital Gonorrhea Caused by Neisseria gonorrhoeae. Antimicrobial Agents and Chemotherapy AAC.01221-18 (2018). doi:10.1128/AAC.01221-18

50. Bell, G. & MacLean, C. The Search for ‘Evolution-Proof’ Antibiotics. Trends in Microbiology 0, (2017).

51. Martínez, J. L., Baquero, F. & Andersson, D. I. Predicting antibiotic resistance. Nat Rev Micro 5, 958–965 (2007).

52. Lázár, V. et al. Antibiotic-resistant bacteria show widespread collateral sensitivity to antimicrobial peptides. Nature Microbiology 3, 718–731 (2018).

53. Baym, M. et al. Spatiotemporal microbial evolution on antibiotic landscapes. Science 353, 1147–1151 (2016).

54. The European Committee on Antimicrobial Susceptibility Testing, E. 2018. EUCAST: Clinical breakpoints. EUCAST: Clinical breakpoints, The European Committee on Antimicrobial Susceptibility Testing Available at: http://www.eucast.org/clinical_breakpoints/. (Accessed: 17th November 2018)

55. Badran, A. H. & Liu, D. R. Development of potent in vivo mutagenesis plasmids with broad mutational spectra. Nat Commun 6, 8425 (2015).

56. O’Neill, A. J. & Chopra, I. Use of Mutator Strains for Characterization of Novel Antimicrobial Agents. Antimicrob Agents Chemother 45, 1599–1600 (2001).

57. Diner, E. J. & Hayes, C. S. Recombineering reveals a diverse collection of ribosomal proteins L4 and L22 that confer resistance to macrolide antibiotics. J Mol Biol 386, 300–315 (2009).

58. Halperin, S. O. et al. CRISPR-guided DNA polymerases enable diversification of all nucleotides in a tunable window. Nature 1 (2018). doi:10.1038/s41586-018-0384-8

59. Garst, A. D. et al. Genome-wide mapping of mutations at single-nucleotide resolution for protein, metabolic and genome engineering. Nat Biotech 35, 48–55 (2017).

60. Nyerges, Á. et al. A highly precise and portable genome engineering method allows comparison of mutational effects across bacterial species. PNAS 201520040 (2016). doi:10.1073/pnas.1520040113

61. Miles, T. J. et al. Novel tricyclics (e.g., GSK945237) as potent inhibitors of bacterial type IIA topoisomerases. Bioorganic & Medicinal Chemistry Letters 26, 2464–2469 (2016).

62. Harkins, C. P. et al. Methicillin-resistant Staphylococcus aureus emerged long before the introduction of methicillin into clinical practice. Genome Biol 18, (2017).

63. Weinstein, Z. B. & Zaman, M. H. Evolution of rifampicin resistance due to substandard drugs in E. coli and M. smegmatis. Antimicrobial Agents and Chemotherapy AAC.01243-18 (2018). doi:10.1128/AAC.01243-18

64. Boeckel, T. P. V. et al. Global antibiotic consumption 2000 to 2010: an analysis of national pharmaceutical sales data. The Lancet Infectious Diseases 14, 742–750 (2014).

65. Taylor, S. N. et al. Single-Dose Zoliflodacin (ETX0914) for Treatment of Urogenital Gonorrhea. New England Journal of Medicine 379, 1835–1845 (2018).

66. Basarab, G. S. et al. Responding to the challenge of untreatable gonorrhea: ETX0914, a first-in-class agent with a distinct mechanism-of-action against bacterial Type II topoisomerases. Scientific Reports 5, 11827 (2015).

67. Foerster, S. et al. Genetic Resistance Determinants, In Vitro Time-Kill Curve Analysis and Pharmacodynamic Functions for the Novel Topoisomerase II Inhibitor ETX0914 (AZD0914) in Neisseria gonorrhoeae. Front. Microbiol. 6, (2015).

68. Damião Gouveia, A. C., Unemo, M. & Jensen, J. S. In vitro activity of zoliflodacin (ETX0914) against macrolide-resistant, fluoroquinolone-resistant and antimicrobial-susceptible Mycoplasma genitalium strains. J Antimicrob Chemother 73, 1291–1294 (2018).

69. The Pew Charitable Trusts. Antibiotics Currently in Clinical Development. Available at: http://www.pewtrusts.org/∼/media/assets/2017/12/antibiotics_currently_in_clinical_development_0_2017.pdf?la=en.9_2017.pdf?la=en. (Accessed: 6th August 2017)

70. Small-Molecule Drug Discovery Suite 2017-4. (Schrödinger, LLC, New York, NY, 2017).

71. Marvin 16.2.1. (Chemaxon (http://www.chemaxon.com), 2016).

72. Harder, E. et al. OPLS3: A Force Field Providing Broad Coverage of Drug-like Small Molecules and Proteins. Journal of Chemical Theory and Computation 12, 281–296 (2016).

73. Berendsen, H. J. C., Postma, J. P. M., van Gunsteren, W. F. & Hermans, J. Interaction Models for Water in Relation to Protein Hydration. in Intermolecular Forces (ed. Pullman, B.) 14, 331–342 (Springer Netherlands, 1981).

74. Kevin J. Bowers, E. C., Huafeng Xu, Ron O. Dror, Michael P. Eastwood, Brent A. Gregersen, John L. Klepeis, Istvan Kolossvary, Mark A. Moraes, Federico D. Sacerdoti, John K. Salmon, Yibing Shan, and David E. Shaw. Scalable Algorithms for Molecular Dynamics Simulations on Commodity Clusters. in (ACM/IEEE, Tampa, Florida, 2006).

75. Schrödinger Release 2017-4: Bioluminate. (Schrödinger, LLC, New York, NY, 2017).

76. Kollman, P. A. et al. Calculating Structures and Free Energies of Complex Molecules:?Combining Molecular Mechanics and Continuum Models. Acc. Chem. Res. 33, 889–897 (2000).

77. Sirin, S., Pearlman, D. A. & Sherman, W. Physics-based enzyme design: Predicting binding affinity and catalytic activity. Proteins: Structure, Function, and Bioinformatics 82, 3397–3409 (2014).

78. Chrencik, J. E. et al. Crystal Structure of Antagonist Bound Human Lysophosphatidic Acid Receptor 1. Cell 161, 1633–1643 (2015).

79. Krishnamurthy, V. R. et al. Glycopeptide analogues of PSGL-1 inhibit P-selectin in vitro and in vivo. Nature Communications 6, 6387 (2015).

80. Lyne, P. D., Lamb, M. L. & Saeh, J. C. Accurate Prediction of the Relative Potencies of Members of a Series of Kinase Inhibitors Using Molecular Docking and MM-GBSA Scoring. J. Med. Chem. 49, 4805–4808 (2006).

81. Greenidge, P. A., Kramer, C., Mozziconacci, J.-C. & Wolf, R. M. MM/GBSA Binding Energy Prediction on the PDBbind Data Set: Successes, Failures, and Directions for Further Improvement. J. Chem. Inf. Model. 53, 201–209 (2013).

82. Ricaurte, D. E. et al. A standardized workflow for surveying recombinases expands bacterial genome-editing capabilities. Microbial Biotechnology 11, 176–188 (2018).

83. Jiang, W., Bikard, D., Cox, D., Zhang, F. & Marraffini, L. A. RNA-guided editing of bacterial genomes using CRISPR-Cas systems. Nat Biotech 31, 233–239 (2013).

84. Umenhoffer, K. et al. Genome-Wide Abolishment of Mobile Genetic Elements Using Genome Shuffling and CRISPR/Cas-Assisted MAGE Allows the Efficient Stabilization of a Bacterial Chassis. ACS Synth. Biol. 6, 1471–1483 (2017).

85. Bódi, Z. et al. Phenotypic heterogeneity promotes adaptive evolution. PLOS Biology 15, e2000644 (2017).

86. Warringer, J., Ericson, E., Fernandez, L., Nerman, O. & Blomberg, A. High-resolution yeast phenomics resolves different physiological features in the saline response. PNAS 100, 15724–15729 (2003).

87. Karcagi, I. et al. Indispensability of Horizontally Transferred Genes and Its Impact on Bacterial Genome Streamlining. Mol Biol Evol 33, 1257–1269 (2016).

88. Dykhuizen, D. E. Experimental Studies of Natural Selection in Bacteria. Annual Review of Ecology and Systematics 21, 373–398 (1990).

89. Altschul, S. F., Gish, W., Miller, W., Myers, E. W. & Lipman, D. J. Basic local alignment search tool. Journal of Molecular Biology 215, 403–410 (1990).

90. DeRyke, C. A., Banevicius, M. A., Fan, H. W. & Nicolau, D. P. Bactericidal Activities of Meropenem and Ertapenem against Extended-Spectrum-β-Lactamase-Producing Escherichia coli and Klebsiella pneumoniae in a Neutropenic Mouse Thigh Model. Antimicrobial Agents and Chemotherapy 51, 1481–1486 (2007).

